# A multiprotein signaling complex sustains AKT and mTOR/S6K activity necessary for the survival of cancer cells undergoing stress

**DOI:** 10.1101/2023.01.03.522657

**Authors:** Oriana Y. Teran Pumar, Matthew R. Zanotelli, Miao-chong Joy Lin, Rebecca R. Schmitt, Kai Su Green, Katherine S. Rojas, Irene Y. Hwang, Richard A. Cerione, Kristin F. Wilson

## Abstract

The ability of cancer cells to survive microenvironmental stresses is critical for tumor progression and metastasis; however, how they survive these challenges is not fully understood. Here, we describe a novel multiprotein complex (DockTOR) essential for the survival of cancer cells under stress, triggered by the GTPase Cdc42 and a signaling partner Dock7, which includes AKT, mTOR, and the mTOR regulators TSC1, TSC2, and Rheb. DockTOR enables cancer cells to maintain a low but critical mTORC2-dependent phosphorylation of AKT during serum deprivation by preventing AKT dephosphorylation through an interaction between phospho-AKT and the Dock7 DHR1 domain. This activity stimulates a Raptor-independent but Rapamycin-sensitive mTOR/S6K activity necessary for survival. These findings address long-standing questions of how Cdc42 signals result in mTOR activation and demonstrate how cancer cells survive conditions when growth factor-dependent activation of mTORC1 is off. Determining how cancer cells survive stress conditions could identify vulnerabilities that lead to new therapeutic strategies.

## Introduction

Cancer cells within a heterogenous tumor and upon their dissemination to secondary tissues during metastatic spread are exposed to a variety of microenvironmental stresses and harsh conditions.^1, 2^ Among these stresses are varying degrees of nutrient, growth factor, and oxygen deprivation as well as resistance to anoikis during detachment from the extracellular matrix that would cause normal healthy cells to succumb to apoptosis. Cancer cells have an exceptional ability to adapt to these conditions and show remarkable tolerance to nutrient deprivation.^3^ As a critical regulator of cell proliferation, growth, and survival, mechanistic target of Rapamycin (mTOR) signaling pathways are often exploited in cancer,^4–8^ and activated mTOR has been shown to support cell survival during growth factor withdrawal mTOR is hyperactivated in most cancers through numerous mechanisms, such as the amplification of receptor tyrosine kinases that signal to phosphatidylinositol-3-kinase (PI3K) (e.g., HER2 and EGFR), gain of function mutations in the catalytic subunit of PI3K (*PIK3CA*), or deletions in PTEN, one of the main negative regulators of the PI3K/mTOR pathway.^9–11^ Upon oncogenic transformation, anabolic processes are thus enhanced and catabolic processes are altered to meet the biosynthetic demands of rapidly growing cancer cells; requirements that are often met by signaling pathways that promote mTOR activity.^12–16^ However, the function and regulation of these pathways in stress conditions such as the absence of growth factors and other nutrients remains largely unknown.

mTOR is a serine/threonine kinase and a member of the PIKK family, which is conventionally known to function within two distinct complexes, mTORC1 and mTORC2.^17–19^ mTORC1 is a Rapamycin-sensitive, nutrient- and mitogen-sensing complex defined by its interaction with the accessory protein Raptor.^20, 21^ Unlike mTORC1, mTORC2 is not inhibited by acute Rapamycin treatment and consists of mTOR, Rictor, mLST8, and mSin1.^22–24^ mTORC2 is activated by multiple mitogenic stimuli and promotes cell survival in response to stresses such as growth factor deprivation.^25, 26^ AKT is a major substrate for mTORC2, which phosphorylates AKT within its hydrophobic motif at Ser473 to enhance its kinase activity. AKT is one of the most activated protein kinases in human cancer^27, 28^ and the mTORC2-AKT axis is widely regarded as a quintessential regulator of apoptosis and cell survival.^29, 30^

mTORC1 is activated by growth factor and amino acid signaling and is recruited to the lysosome by the interaction between Raptor and the heterodimeric GTPase RagA/B-C/D complex, which is activated by the Ragulator-v-ATPase.^31–33^ When bound to the lysosome, mTORC1 interacts with its direct activator, the small GTPase Ras homolog enriched in brain (Rheb).^34^ Rheb stimulates mTORC1 kinase activity, triggering anabolic processes, including protein and lipid synthesis, while inhibiting catabolic processes such as autophagy. AKT plays an essential role in how growth factors activate mTORC1 by catalyzing the phosphorylation of Tuberous Sclerosis Complex 2 (TSC2), which disables its ability to act together with TSC1 to negatively regulate mTOR activation by serving as a Rheb GTPase Activating Protein (GAP).^35^ While Rheb and the Rags have well established roles in activating mTORC1, members of the Rho GTPase family, traditionally known for their involvement in cytoskeletal remodeling,^36^ have also been implicated in regulating mTOR activity and in driving tumorigenesis.^37–39^ The GTPase Cdc42 has been shown to signal to mTOR and thus promote the phosphorylation of its downstream target the p70 S6 kinase (S6K).^40–44^ Thus far, the mechanism responsible for how Cdc42 signals mTOR activation has not yet been elucidated.

Here, we identify a multiprotein complex, referred to as DockTOR, that includes Cdc42 and the Cdc42/Rac GEF and Cdc42-signaling effector, Dock7, that can protect and maintain low-levels of AKT activity as well as promote a growth factor-independent activation of mTOR/S6K during cellular stress. We show that within DockTOR, Dock7 serves as a binding partner for Cdc42, AKT, mTOR, the mTORC1-associated proteins TSC1/2, and Rheb. Stresses, such as serum deprivation, promote the assembly of a complex that stimulates a pro-survival signaling activity by sustaining a low-level of AKT phosphorylation at the mTORC2 phospho-site Ser473, which then promotes a Rheb-dependent but Raptor-independent activation of mTOR and subsequent S6K phosphorylation. Dock7 belongs to the atypical Dock180 family of Rho GEFs with two conserved domains, Dock Homology Region 1 (DHR1) and DHR2.^45, 46^ DHR2 contains a GEF domain that can activate either Cdc42 or Rac1, an allosteric binding site for activated Cdc42, and a putative dimerization region, while DHR1 has a C2-like motif ascribed to membrane phospholipid binding in other Dock proteins but with no reported signaling function in Dock7.^47, 48^ We find that the binding of activated Cdc42 to an allosteric site within DHR2, the dimerization of Dock7 through the DHR2 domains, and the function of the C2-like motif of DHR1, are each necessary for DockTOR to sustain AKT activation and stimulate mTOR/S6K activity during stress conditions and promote survival in cancer cells.

## Results

### Cdc42 works together with its signaling partner Dock7 to activate mTOR

It has been known for some time that Cdc42 is able to stimulate mTOR activity with important consequences in various biological contexts.^40–44^ Ectopic expression of the constitutively active form of Cdc42 (Cdc42 Q61L) in HeLa cells stimulated S6K phosphorylation at Thre389 (p-S6K), a frequently used read-out for mTOR stimulation^49^, to a comparable degree as Heregulin (an activator of ErbB2/HER2) or upon ectopic expression of the small GTPase Rheb, which directly activates mTORC1^34^ (Figure 1A). A clue regarding how Cdc42 signals to mTOR came from an earlier report suggesting that Dock7, a member of the Dock180 family of Rho GEFs, that we have shown to be both a GEF and an effector for Cdc42,^50^ is capable of interacting with the Rheb GAP complex TSC1/2.^51^

**Figure 1.**
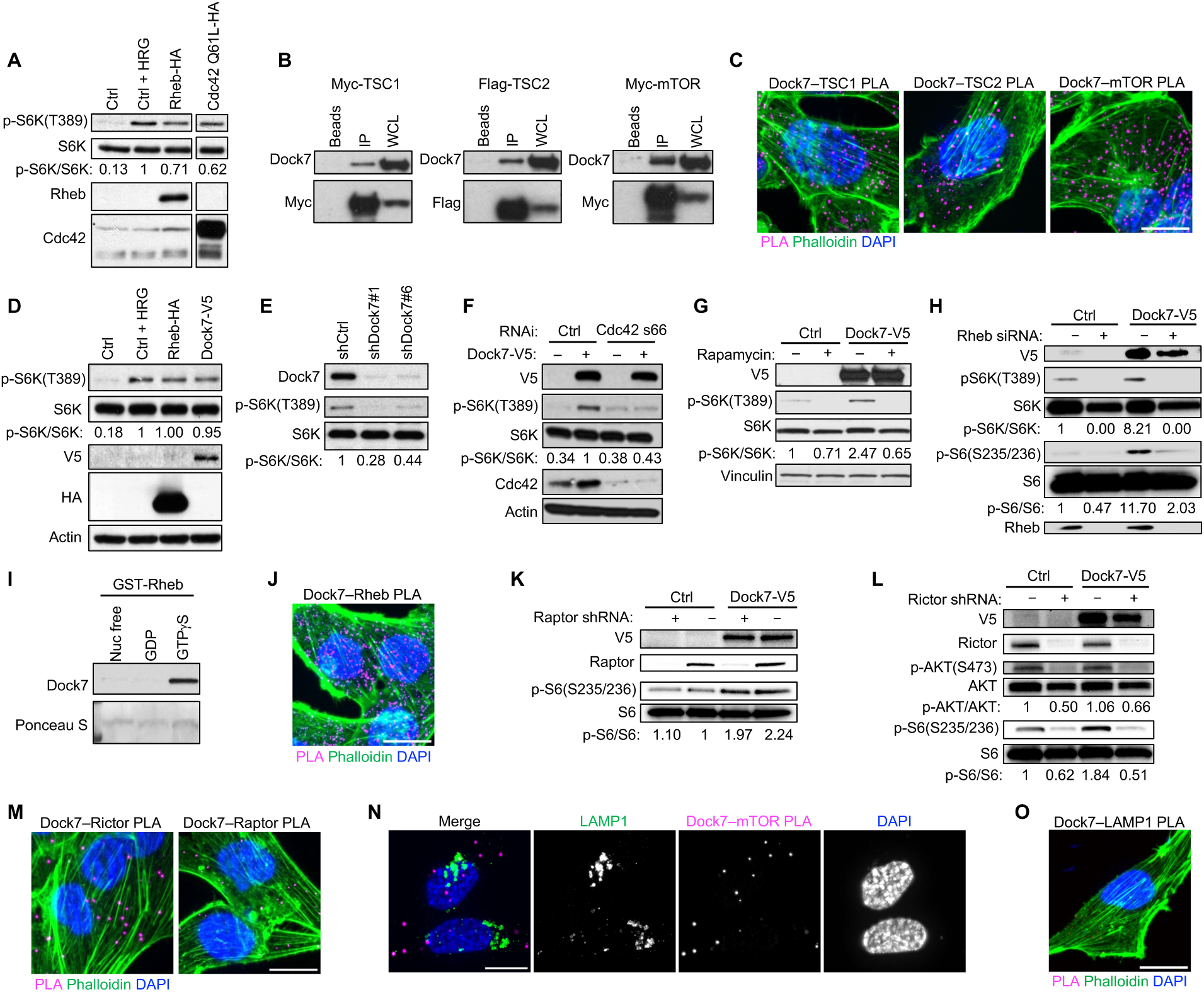
Dock7 interacts with TSC1, TSC2, Rheb, and mTOR and stimulates S6K in a Cdc42-dependent manner that requires Rictor, but not Raptor. (**A**) Western blot of serum-starved HeLa cells transiently transfected with Rheb-HA or Cdc42 Q61L-HA. Heregulin (HRG) treatmen was used as a control to confirm mTORC1 signaling. (**B**) Western blot of endogenous Dock7 co-immunoprecipitating with ectopically expressed Flag-TSC2, Myc-TSC1, or Myc-mTOR from HEK-293T cells. (**C**) PLA of interactions between endogenous Dock7 and TSC1, TSC2, or mTOR in MDA-MB-231 in complete media. (**D**) Western blot of S6K activity in HeLa cells transiently transfected with either Dock7 or Rheb and starved overnight and treated with HRG as a positive control. (**E**) Western blot of S6K activity in serum-starved HeLa cells following Dock7 knockdown. (**F**) Western blot of S6K activity in serum-starved HeLa cells with Dock7-V5 transiently overexpressed, and Cdc42 knocked down using siRNA. (**G**) Western Blot of S6K activity in serum-starved HeLa cells transiently expressing Dock7-V5 in presence of either a vehicle control or Rapamycin. (**H**) Western blot of S6K or S6 phosphorylation in serum-starved MBA-MD-231 cells transiently overexpressing Dock7-V5 in the presence or absence Rheb knockdown using siRNA. (**I**) Recombinant GST-Rheb proteins loaded with either GTPγS, GDP, or EDTA-treated were used to pull-down endogenous Dock7 from HEK-293T cells. Ponceau S was used to visualize the GST proteins. (**J**) PLA of Dock7–Rheb interactions in MDA-MB-231 cells grown in serum-free media. (**K**) Western blot of S6K phosphorylation in MDA-MB-231 cells following semi-stable Raptor knockdown using shRNA and Dock7-V5 overexpression. (**L**) Western blot of AKT and S6K phosphorylation in MDA-MB-231 cells following semi-stable Rictor knockdown using shRNA and Dock7-V5 overexpression. (**M**) PLA of Dock7–Rictor and Dock7–Raptor interactions in MDA-MB-231 cells in serum-free media. (**N**) PLA of Dock7– mTOR interactions with immunofluorescence staining of LAMP1 in MDA-MB-231 cells in serum-free media. (**O**) PLA of Dock7–LAMP1 interactions in MDA-MB-231 cells in serum-free media. Scale bar = 15 μm.

To establish that Dock7 can in fact interact with TSC1/2, we transiently transfected HEK-293T cells with either epitope-tagged TSC1 or TSC2 and isolated these proteins via immunoprecipitation. Co-immunoprecipitation (co-IP) experiments showed that endogenous Dock7 associated with both Myc-tagged TSC1 and Flag-tagged TSC2. We also found that Myc-tagged mTOR was capable of co-immunoprecipitating endogenous Dock7 (Figure 1B). Consistent with these observations, Dock7 co-migrated with mTOR, TSC1, and TSC2 as a high molecular weight species upon performing Blue Native-PAGE (BN-PAGE) using HEK-293T cell lysates,^52^ a technique used to characterize the components of large complexes based on their native molecular mass (Figure S1). As a complementary approach, we used proximity ligation assays (PLA), which can quantitatively visualize proteins that are within tens of nanometers of each other as fluorescent puncta *in situ.*^53, 54^ PLA in MDA-MB-231 highly metastatic breast cancer cells yielded robust puncta and corroborated the results obtained by immunoprecipitation indicating interactions between endogenous Dock7 and TSC1, TSC2, and mTOR (Figure 1C).

Given that Dock7 associated with mTORC1 regulators TSC1/2 as well as mTOR, we examined its ability to stimulate S6K activity. Like activated Cdc42, the ectopic expression of Dock7 promoted S6K phosphorylation during serum starvation, while knocking down Dock7 reduced basal levels of S6K/S6 phosphorylation during serum starvation (Figures 1D and 1E). Notably, Cdc42 was required for Dock7 to activate S6K, as Dock7 overexpression was not able to stimulate S6K phosphorylation following Cdc42 knockdown (Figure 1F). These observations identify a previously unknown role for Dock7 in the Cdc42-dependent activation of mTOR.

### Dock7 stimulation of mTOR/S6K activity requires both mTORC2 and Rheb but not Raptor

We next sought to further define the mechanism by which Dock7 mediates the activation of mTOR in nutrient-deprived cells. Treatment with Rapamycin, an mTORC1-specific inhibitor, blocked the ability of ectopically expressed V5-tagged Dock7 to promote the phosphorylation of S6K (Figure 1G), and knocking down Rheb prevented Dock7 from stimulating the phosphorylation of both S6K and its major target ribosomal S6 (Figure 1H). Rheb, specifically when in an activated state (i.e., bound to the GTP analog GTPγS), was capable of associating with Dock7 in pull-down experiments (Figure 1I). GST pull-down assays were supported by PLA, which showed a substantial number of signals between endogenous Dock7 and Rheb during serum starvation (Figure 1J).

Given the ability of Dock7 to activate S6K in serum-starved cells, we examined the role of the mTORC1-defining subunit, Raptor, in this process. Unexpectedly, when Raptor was knocked down there was no effect on the activation of S6K, as read out by S6 phosphorylation, in either serum-starved control cells that endogenously express Dock7 or Dock7 over-expressing cells (Figure 1K). We next looked at the participation of Rictor, the mTORC2 defining subunit, for its ability to mediate Dock7 stress-dependent signaling. As expected, the Rictor knockdown resulted in reductions in AKT phosphorylation in both mock and Dock7 over-expressing cells; moreover, it also reduced the phosphorylation of S6 (Figure 1L). S6K is not a direct substrate of mTORC2, therefore, the Rictor-dependent activation of S6K suggests Dock7 acts downstream of mTORC2. We also observed relatively strong PLA signal between endogenous Dock7 and Rictor, compared to Dock7 and Raptor in serum-starved MDA-MB-231 cells (Figure 1M). Immunofluorescence staining failed to show significant co-localization of Dock7 with the endosomal-lysosome marker LAMP1 during serum deprivation and the Dock7–mTOR PLA signal similarly did not localize with LAMP1 (Figures 1N and S2). PLA also failed to show proximity between Dock7 and LAMP1, suggesting that Dock7 stimulates S6K activity independently of lysosomal involvement (Figure 1O).

### Dock7, together with mTOR, AKT and S6K, promotes signaling activities essential for cancer cell survival

Searching the Cancer Genome Atlas (TCGA) Breast Cancer Dataset (BRCA), we found that unlike other members of the Dock-C subfamily of Dock180 proteins, Dock7 is highly upregulated in triple-negative breast cancer patients (Figures 2A and 2B). Examining Dock7 protein expression levels among a panel of 10 different breast cancer cell lines, half of which were HER2^+^, ER^+^/PR^+^/HER2^+^, or ER^+^/PR^+^ cell lines, while the remaining half were triple-negative breast cancer cell lines, also showed that triple-negative breast cancer cells exhibited higher Dock7 protein expression (Figure 2C). We therefore investigated the effects of Dock7 on the ability of breast cancer cells to undergo anchorage-independent growth, a hallmark of transformation and a cellular stressor as cells are forced to grow in the absence of a substratum.^55^ Upon knocking down Dock7, we observed a striking decrease in the ability of triple-negative MDA-MB-231 breast cancer cells, as well as receptor-positive SK-BR-3 and MCF7 breast cancer cells, to form colonies in soft agar (Figure 2D). The knockdown of Dock7 also impacted the ability of HeLa cervical cancer cells and A549 lung cancer cells to form colonies (Figure 2E and 2F), establishing that Dock7 is necessary for the transformed properties of multiple cancer cell types.

**Figure 2.**
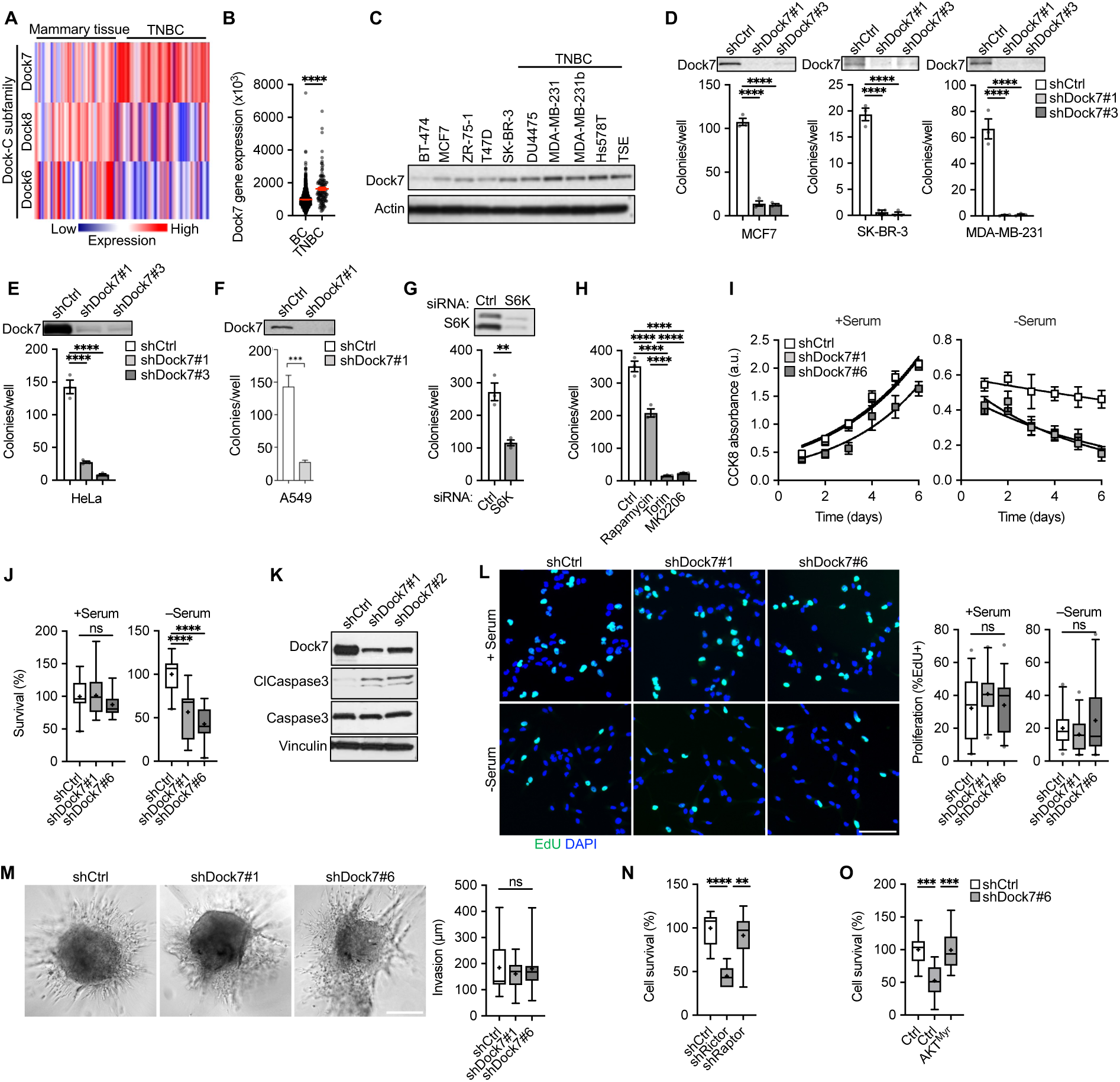
Dock7 is required for the transformation and survival of cancer cells and suggests functional roles for AKT, mTOR and S6K. (**A**) The Cancer Genome Atlas (TGCA) expression profile for Dock-C family members in triple-negative breast cancer (TNBC) and mammary tissue. (**B**) Dock7 mRNA levels across patients with either receptor-positive (BC) or TNBC. (**C**) Western blot of Dock7 protein expression levels across breast cancer cell lines. (**D**) Soft agar colony formation of MDA-MB-231, SK-RB-3 and MCF7 breast cancer cells following knockdown of Dock7. (**E**) Soft agar colony formation of HeLa cells after knockdown of Dock7. (**F**) Soft agar colony formation of A549 cells after knockdown of Dock7. (**G**) Soft agar colony formation of HeLa cells following S6K knockdown using siRNA. (**H**) Soft agar colony formation of HeLa cells grown in complete media containing either vehicle control, Rapamycin, Torin, or MK2206. (**I**) CCK8 absorbance over 6 days in Dock7 knockdown MDA-MB-231 cells in the presence or absence of serum. Lines show exponential growth curves. (**J**) Cell survival of MDA-MB-231 cells following Dock7 knockdown after 2 days in the presence or absence of serum. (**K**) Western blot of Cleaved Caspase-3 in serum-starved HeLa cells after Dock7 knockdown. (**L**) EdU proliferation assays performed in the presence or absence of serum for MDA-MB-231 cells following Dock7 knockdown. (**M**) MDA-MB-231 breast cancer cell spheroid invasion in collagen over 24-48 h post-embedding, following Dock7 knockdown (**N**) Cell survival of MDA-MB-231 cells during serum starvation following Rictor and Raptor knockdown. (**O**) Cell survival of MDA-MB-231 cells during serum starvation following Dock7 knockdown and overexpression of a constitutively active AKT mutant construct (AKT^Myr^). Data shown as median ± interquartile range (box), 5^th^– 95^th^ percentiles (whiskers), and mean (+), or mean ± s.d. **P<0.01, ***P<0.001, ****P<0.0001, ns = not significant. Scale bar = (**L**) 100 μm and (**M**) 500 μm.

We then examined whether the relationships observed between Dock7, mTOR and S6K, might be important for the role of Dock7 in transformation. Both the knockdown of S6K and treatment of HeLa cells with Rapamycin resulted in an approximately 50% decrease in soft agar colony formation (Figures 2G and 2H). Consistent with Rictor-associated mTOR activity being important in Dock7 signaling, we found that treating HeLa cells with Torin, a pan mTOR inhibitor, or MK2206, an AKT inhibitor, fully blocked growth in soft agar (Figure 2H). The fact that knocking down S6K, and Rapamycin treatment, partially blocked transformation is likely due to AKT triggering survival and anti-apoptotic activities that are additional to the activation of an mTORC1-like activity.

Given that AKT promotes cell survival, proliferation, and invasion^56^ and is a negative regulator of apoptosis,^57, 58^ we next determined the role of Dock7 in each of these transformed phenotypes. MDA-MB-231 cells showed a decrease in cell viability following Dock7 knockdown during serum-free conditions, compared to complete media, as read-out using a CCK8 assay kit, as well as when directly counting cell numbers (Figures 2I and 2J). Apoptosis, as measured by increases in the levels of Cleaved Caspase-3, was elevated in serum-starved HeLa cells when Dock7 was knocked down (Figure 2K). In contrast, Dock7 knockdown did not affect cell proliferation regardless of culture conditions (Figure 2L). The expression of Dock7 was also dispensable for the invasive activity of MDA-MB-231 cells, as measured by spheroid invasion in 3D collagen (Figure 2M). Cell survival during serum starvation required mTORC2 signaling, as Rictor knockdown reduced survival while Raptor knockdown had no effect (Figure 2N). Additionally, Dock7-dependent loss of cell survival during serum deprivation could be rescued by ectopically expressing a myristoylated, constitutively active form of AKT (AKT^Myr^) (Figure 2O). Together, these findings suggest that Dock7, while not required for mitogenic growth, plays a role in regulating AKT activity necessary for cells to survive nutrient deprivation and resist anoikis.

### Dock7 protects AKT from dephosphorylation at Ser473

We then examined the ability of Dock7 to interact with AKT. Co-IP experiments utilizing the co-expression of V5-tagged Dock7 and Flag-tagged AKT in growing HEK-293T cells showed that a relatively small, but detectable amount of transiently expressed AKT co-precipitated with Dock7-V5 (Figure 3A). We used PLA in MDA-MB-231 cells to confirm that Dock7 was in sufficient proximity to interact with AKT *in situ*. While PLA showed a similar number of interactions between Dock7 and AKT regardless of culture conditions, the PLA signal between Dock7 and phospho-AKT(S473) increased upon serum deprivation, suggesting that Dock7-bound AKT becomes phosphorylated by mTORC2 under these conditions (Figures 3B and 3C). Dock7 also appeared to help maintain the interactions between phosphorylated/activated AKT and its target, TSC2, as knocking down Dock7 significantly reduced the PLA signal between phospho-AKT(S473) and phospho-TSC2(T1462) (Figure 3D).

**Figure 3.**
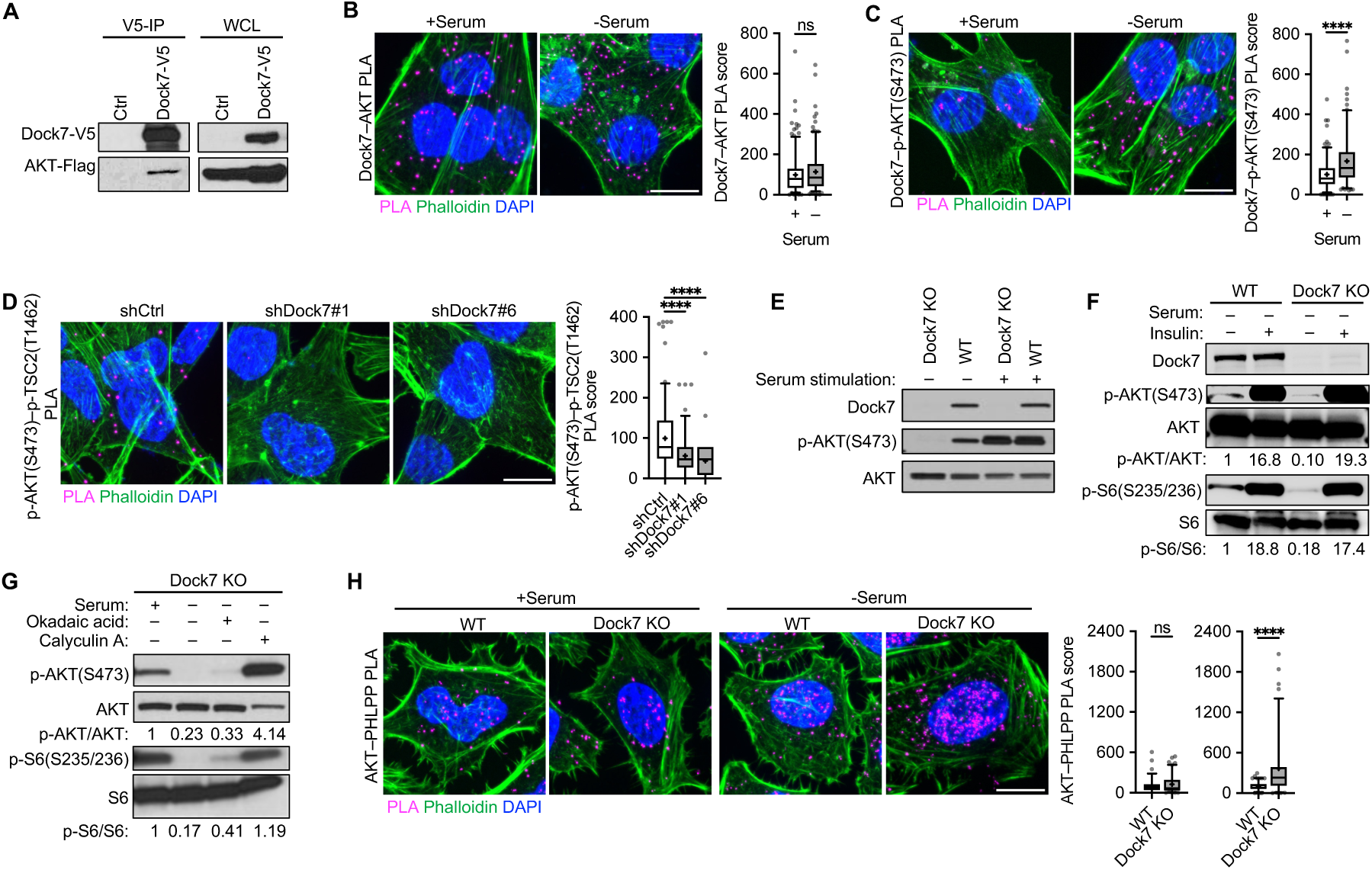
Dock7 protects AKT from dephosphorylation to enhance cancer cell survival during serum deprivation. (**A**) Immunoprecipitation of transiently overexpressed Flag-tagged AKT and Dock7-V5 in HEK-293T cells resolved on an SDS-PAGE gel for Western analysis. (**B**, **C**) PLA image and PLA score of (**B**) Dock7–AKT and (**C**) Dock7–p-AKT(S473) interactions in MDA-MB-231 cells in complete media or following overnight serum starvation. (**D**) p-AKT(S473)–p-TSC2(T1462) PLA image and PLA score in MDA-MB-231 cells following Dock7 knockdown and overnight serum starvation. (**E**) HeLa cells were seeded and allowed to recover for a day before changing media to serum-free media for 24 h then serum stimulation (10% FBS) was performed with dialyzed serum for 1 h and Western blotting of p-AKT(S473) was performed. (**F**) HeLa cells were seeded and allowed to recover for a day before changing media to serum-free media for 24 h. Cells were treated with and without 100 nM insulin for 1 h before being collected and used for Western blot of p-AKT(S473) or p-S6(S235/236). (**G**) Western blot analysis to indicate the phosphorylation status of AKT and S6 in Crispr-Cas9 Dock7 KO HeLa cells following treatment with phosphatase inhibitors Okadaic acid and Calyculin A. (**H**) PLA image and score of AKT–PHLPP interactions in Dock7 KO and WT HeLa cells in the presence and absence of serum. Data shown as median ± interquartile range (box), 5^th^–95^th^ percentiles (whiskers), and mean (+). ****P<0.0001, ns = not significant. Scale bar = 15 μm.

We then created a Dock7 knock-out (KO) model using Crispr-Cas9. We found that the genetic ablation of Dock7 was lethal to MDA-MB-231 cells; however, we were able to generate a Dock7 KO HeLa cell line which showed a dramatic reduction in soft agar colony formation, similar to the results we observed in knockdown experiments (Figures S3A and S3B). Like the case when knocking down Dock7, Dock7 KO cells were unable to survive serum-free conditions relative to wild-type (WT) cells and displayed enhanced rates of apoptosis (Figures S3C and S3D). Proliferation was unaffected by the Dock7 KO (Figure S3E). Murine Embryonic Fibroblasts (MEFs) derived from Dock7 KO mice did not experience a decrease in survival when grown in serum-free conditions suggesting that the ability of Dock7 to provide a survival benefit may be specific for cancer cells (Figures S3F and S3G).

In both Dock7 KO and WT cells, serum- and insulin-stimulated AKT phosphorylation at Ser473, as well as the phosphorylation of the S6K substrate S6 at Ser235/236, to the same robust extents (Figures 3E and 3F). The basal phosphorylation levels of both AKT and S6, however, were diminished specifically in serum-starved Dock7 KO cells. Dock7 KO cells were then cultured in either full-serum or serum-free media overnight and treated with Okadaic Acid or Calyculin A, inhibitors of serine/threonine protein phosphatases.^59^ A slight rescue of signaling activities was observed following Okadaic Acid treatment, but upon inhibition of phosphatase activity with the more potent inhibitor, Calyculin A, the phosphorylation of AKT at Ser473 and that of its downstream effector S6, were maintained in Dock7 KO cells during serum deprivation (Figure 3G). We further probed potential interactions between AKT and PHLPP, a phosphatase that shows substrate specificity for AKT phosphorylation at Ser473,^60^ by PLA. There were significant changes in the AKT–PHLPP PLA signal in Dock7 KO cells compared to WT cells when cultured in serum-free media, but not under normal growth conditions (Figure 3H). Together, these data suggest that Dock7 preserves AKT phosphorylation and its activated state by protecting it from dephosphorylation.

### Both evolutionarily conserved Dock domains, DHR1 and DHR2, contribute to maintaining AKT activation and the transforming potential of Dock7

To better understand how Dock7 interacts with AKT, we assessed the roles of the two conserved Dock7 domains (Figure 4A). For these studies, we used constructs encoding a DHR2 limit domain (DHR2) and a DHR1 domain with a C-terminal 183 amino acid addition of the downstream linker region (DHR1L) that was anticipated to express better than the DHR1 limit domain (DHR1E for DHR1 exact) (Figure S4A). Compared to DHR1E, DHR1L showed a greater ability to interact with AKT and phosphorylate AKT as well as reverse the increase in AKT–PHLPP interactions observed when Dock7 is knocked down (Figures S4B-D). DHR1L was relatively also more effective than DHR1E in rescuing cell survival following Dock7 knockdown (Figure S4E), suggesting that the linker region of Dock7 may play an important role in Dock7 interactions with AKT and its downstream signaling partners.

**Figure 4.**
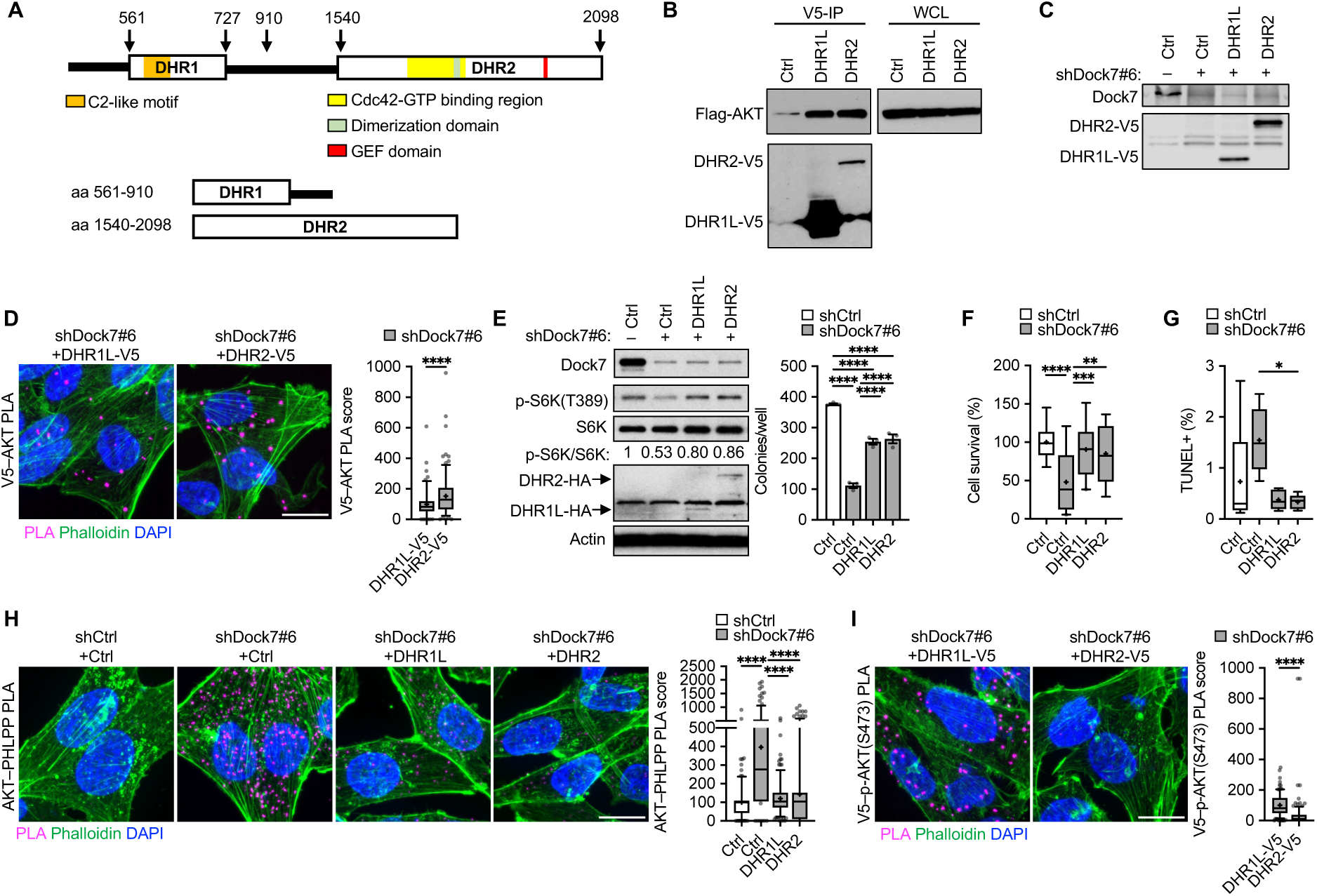
Dock7 DHR1 and DHR2 domains both interact with AKT and provide phenotypic rescue after loss of Dock7, while DHR1 protects AKT from dephosphorylation. (**A**) Schematic of Dock7 and its features, and Dock7 limit domain constructs. (**B**) Flag-tagged AKT was transiently overexpressed in HEK-293T cells semi-stably expressing either DHR1-V5 or DHR2-V5. Immunoprecipitation was performed using anti-V5 beads and complexes were resolved on an SDS-PAGE gel for Western analysis. (**C**) Western blot of DHR1L-V5 and DHR2-V5 expression after Dock7 knockdown in MDA-MB-231 cells. (**D**) V5–AKT PLA image and score following Dock7 knockdown and DHR1 and DHR2 overexpression in MDA-MB-231 cells after overnight serum starvation. (**E**) Western blot (left panel) and soft agar colony formation (right panel) of HeLa cells with Dock7 knocked down and DHR1 and DHR2 overexpressed. (**F**) Cell survival of MDA-MB-231 cells following Dock7 knockdown and DHR1 and DHR2 overexpression after 2 days in serum-free media. (**G**) TUNEL assay of MDA-MB-231 cells following Dock7 knockdown and DHR1 or DHR2 overexpression after overnight serum starvation. (**H**, **I**) PLA image and score of (**H**) AKT–PHLPP and (**I**) V5–AKT interactions in serum-starved MDA-MB-231 following Dock7 knockdown and V5-tagged limit domain overexpression. Data shown as median ± interquartile range (box), 5^th^–95^th^ percentiles (whiskers), and mean (+), or mean ± s.d. *P<0.05, **P<0.01, ***P<0.001, ****P<0.0001, ns = not significant. Scale bar = 15 μm.

Co-immunoprecipitated protein complexes isolated from HEK-293T cells semi-stably expressing V5-tagged DHR1L or DHR2 and ectopically expressing Flag-AKT showed the ability of both DHR1L and DHR2 to interact with AKT (Figure 4B). Given that significantly less DHR2 was expressed in these cells compared to DHR1L, their comparable effectiveness in co-immunoprecipitating with AKT suggests that AKT undergoes a stronger interaction with the DHR2 domain. In MDA-MB-231 Dock7 knockdown cells with each domain overexpressed, PLA similarly showed increased interactions between DHR2 and endogenous AKT compared to DHR1L and AKT during serum deprivation (Figures 4C and 4D). Low-levels of ectopically expressed Dock7 domains in HeLa cells depleted of endogenous Dock7 partially restored mTOR activity, as read-out by the phosphorylation of S6K at Thre389, and soft-agar colony formation (Figure 4E), as well as the ability of MDA-MB-231 cells to survive serum deprivation and to be protected from undergoing apoptosis (Figures 4F and 4G). Both DHR1L and DHR2 showed an ability to associate with AKT and decrease AKT–PHLPP interactions (Figure 4H). However, PLA signal between p-AKT(S473) and DHR1L expressed in Dock7 knockdown cells were increased compared to DHR2 (Figure 4I), suggesting the DHR1 domain may be primarily responsible for protecting phosphorylated AKT from dephosphorylation.

### Examining the unique role of the DHR1 domain of Dock7

To date, a specific function for the DHR1 domain of Dock7 has not been defined, so we sought to identify its role in the ability of Dock7 to maintain AKT phosphorylation and activate mTOR during serum deprivation. Given DHR1 contains a putative C2-like motif that has been shown to bind phospholipids and to be important in the subcellular localization of Dock180 and Dock2,^47, 61^ we focused our attention on its possible role in maintaining AKT phosphorylation. The sequences of the eleven Dock180-family proteins were aligned and two conserved positive amino acid residues on Dock7 were identified that might be capable of mediating interactions with a negatively charged binding partner. Site-directed mutagenesis was performed to create a DHR1L C2 mutant (DHR1L-C2M) by substituting an alanine for each of the two conserved arginine residues (R576A, R581A).

Although PLA suggested no differences between DHR1L constructs in the number of interactions with AKT (Figure 5A), ectopic expression of DHR1L showed increased PLA signal with p-AKT(S473) relative to DHR1L-C2M in serum-starved MDA-MB-231 cells (Figure 5B). Similarly, while both the expression of DHR1L and DHR2 were able to reverse the interactions between AKT and PHLPP that occur upon the knockdown of Dock7, the expression of DHR1L-C2M appeared to cause a further increase in these interactions compared to knocking down Dock7 alone (Figure 5C). Survival assays in Dock7 knockdown cells with V5-tagged DHR1L, DHR1L-C2M, and DHR2 domains versus a vector control ectopically expressed to create semi-stable cell lines, showed that both DHR1L and DHR2 partially restored cell survival and prevented apoptosis under serum-free conditions, whereas DHR1L-C2M was significantly less effective (Figures 5D and 5E). We then examined the role of the C2-like motif within the context of full-length Dock7. We first compared the ability of V5-tagged full-length WT Dock7 (Dock7-WT-V5) versus Dock7-C2M-V5 to interact with p-AKT(S473) in serum-starved MDA-MB-231 cells (Figure 5F). Like the DHR1 limit domain construct containing this mutation, Dock7-C2M-V5 showed a marked decrease in the V5–p-AKT(S473) PLA signal compared to Dock7-WT-V5 (Figure 5G), consistent with the suggestion that Dock7, via the C2-like motif of DHR1, preferentially engages phosphorylated AKT during serum deprivation. AKT–PHLPP PLA signal was also significantly increased in Dock7-C2M cells relative to control and Dock7-WT (Figure 5H).

**Figure 5.**
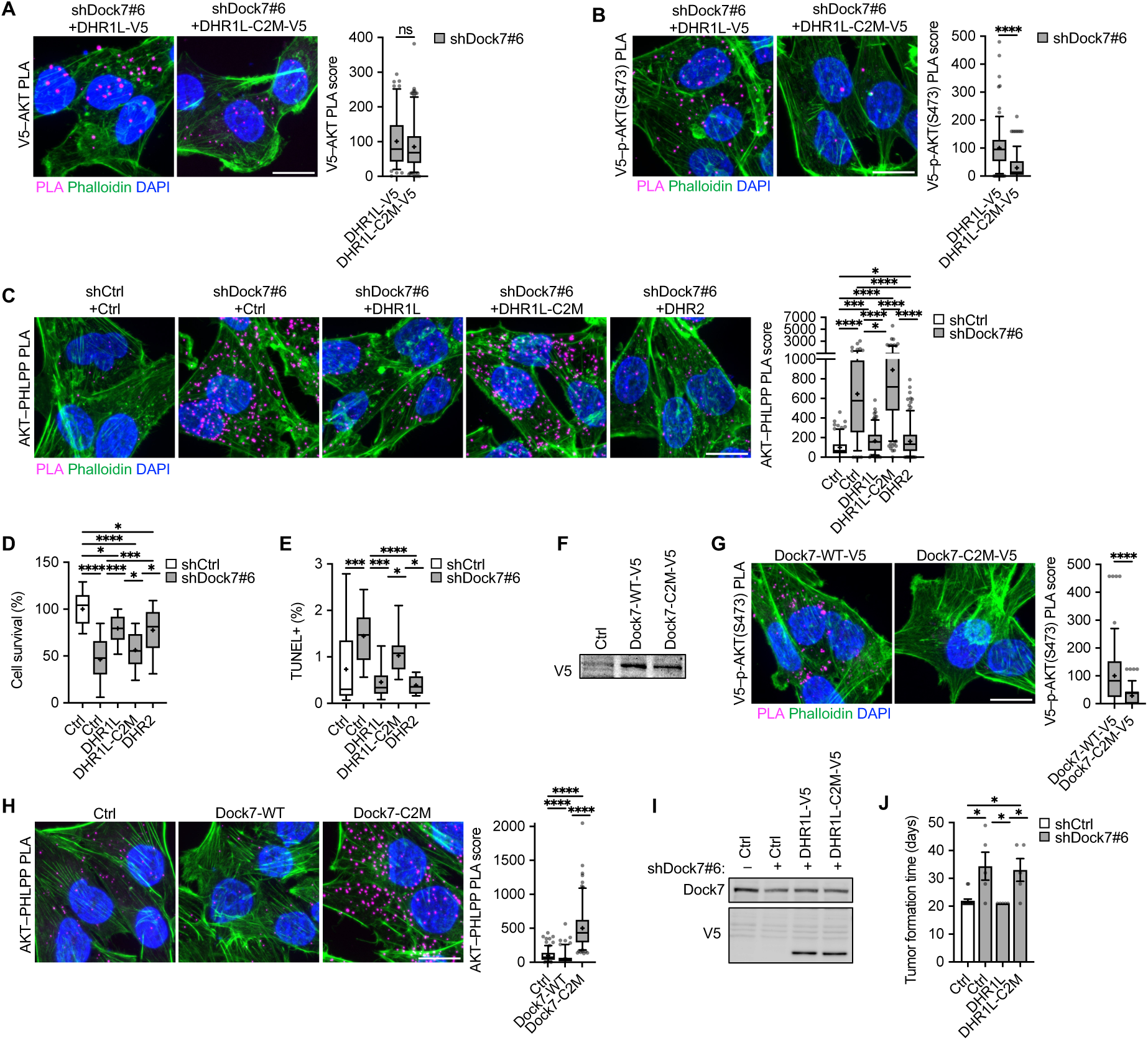
The DHR1 C2-like motif is necessary to protect phosphorylated AKT and sustain AKT and mTOR/S6K activity. (**A**-**C**) PLA image and score of (**A**) V5–AKT, (**B**) V5–p-AKT(S473), and (**C**) AKT–PHLPP interactions in serum-starved MDA-MB-231 following Dock7 knockdown and DHR1,DHR1-C2M and DHR2 overexpression. (**D**) MDA-MB-231 cell survival following Dock7 knockdown and overexpression of DHR constructs. (**E**) TUNEL assay of MDA-MB-231 cells following Dock7 knockdown and DHR limit domain constructs overexpression after overnight serum starvation. (**F**) Western blot of Dock7-V5 and Dock7-C2M-V5 in MDA-MB-231 cells. (**G, H**) PLA image and score of (**G**) V5–p-AKT(S473) and (**H**) AKT–PHLPP interactions in MDA-MB-231 cells transiently overexpressing WT Dock7 and Dock7-C2M after overnight serum starvation. (**I**, **J**) NSG mouse tumor xenograft. (**I**) MDA-MB-231 cells following Dock7 knockdown and DHR1L and DHR1L-C2M limit domain overexpression used for xenografts. (**J**) Tumor formation time. Data shown as median ± interquartile range (box), 5^th^–95^th^ percentiles (whiskers), and mean (+), or mean ± s.d. *P<0.05, ***P<0.001, ****P<0.0001, ns = not significant. Scale bar = 15 μm.

We then performed mouse xenograft experiments to examine the role of Dock7 and the unique actions of its DHR1 domain in survival *in vivo* during tumor formation by implanting MDA-MB-231 (vector control) cells, Dock7 knockdown cells, and Dock7 knockdown cells in which either DHR1L or DHR1L-C2M were semi-stably expressed (Figure 5I). As expected, Dock7 knockdown cells were not able to form tumors compared to the control MDA-MB-231 cells (Figure 5J). Interestingly, the expression of DHR1L in Dock7 knockdown MDA-MB-231 breast cancer cells helped the cells to survive the initial implantation and form tumors, while cells expressing DHR1-C2M were less effective at forming tumors (Figures 5J and S5).

### The DHR2 domain functions as a dimerization-dependent Cdc42 effector to promote AKT-mTOR activity

We next focused on the DHR2 domain of Dock7 where Cdc42 interactions occur to better understand how Cdc42 works with Dock7 to promote AKT and mTOR/S6K activation. GST pull-down experiments showed the ability of GTPγS-bound forms of Cdc42 to associate with Dock7 indicating that Dock7 can act as a Cdc42 signaling effector as well as a GEF^50^ (Figure 6A), and we observed that Cdc42 interactions with endogenous Dock7 are enhanced in the absence of serum (Figure 6B). Given the most notable feature of DHR2 domains is their ability to promote guanine nucleotide exchange, we examined a mutant that contains a substitution of an alanine for the conserved, catalytic Val1991 residue within the GEF domain that is known to render it GEF-defective^62^ (GEF-defective mutant, GDM). To test the importance of the Dock7 GEF activity in Dock7 interactions with phosphorylated AKT, we compared the ability of the V5-tag of Dock7-WT-V5 and Dock7-GDM-V5 to associate with p-AKT(S473) in serum-starved cells by PLA. These experiments showed no difference in V5–p-AKT(S473) PLA signal between Dock7-WT-V5 and Dock7-GDM-V5, suggesting that the Cdc42-GEF activity of Dock7 was not necessary for it to interact with active AKT during serum deprivation (Figure 6C). Overexpression of Dock7-GDM also promoted AKT and S6K phosphorylation to a similar extent as Dock7-WT during serum starvation, indicating that under these conditions the GEF activity of Dock7 is dispensable for its ability to activate mTOR. However, Dock7-GDM still required Cdc42 to stimulate AKT and S6K (Figure 6D). Collectively, these results suggest that the binding of GTP-bound Cdc42 to the allosteric site located N-terminal to the GEF domain and proximal to a putative dimerization domain may be necessary to promote these signaling activities.^50^

**Figure 6.**
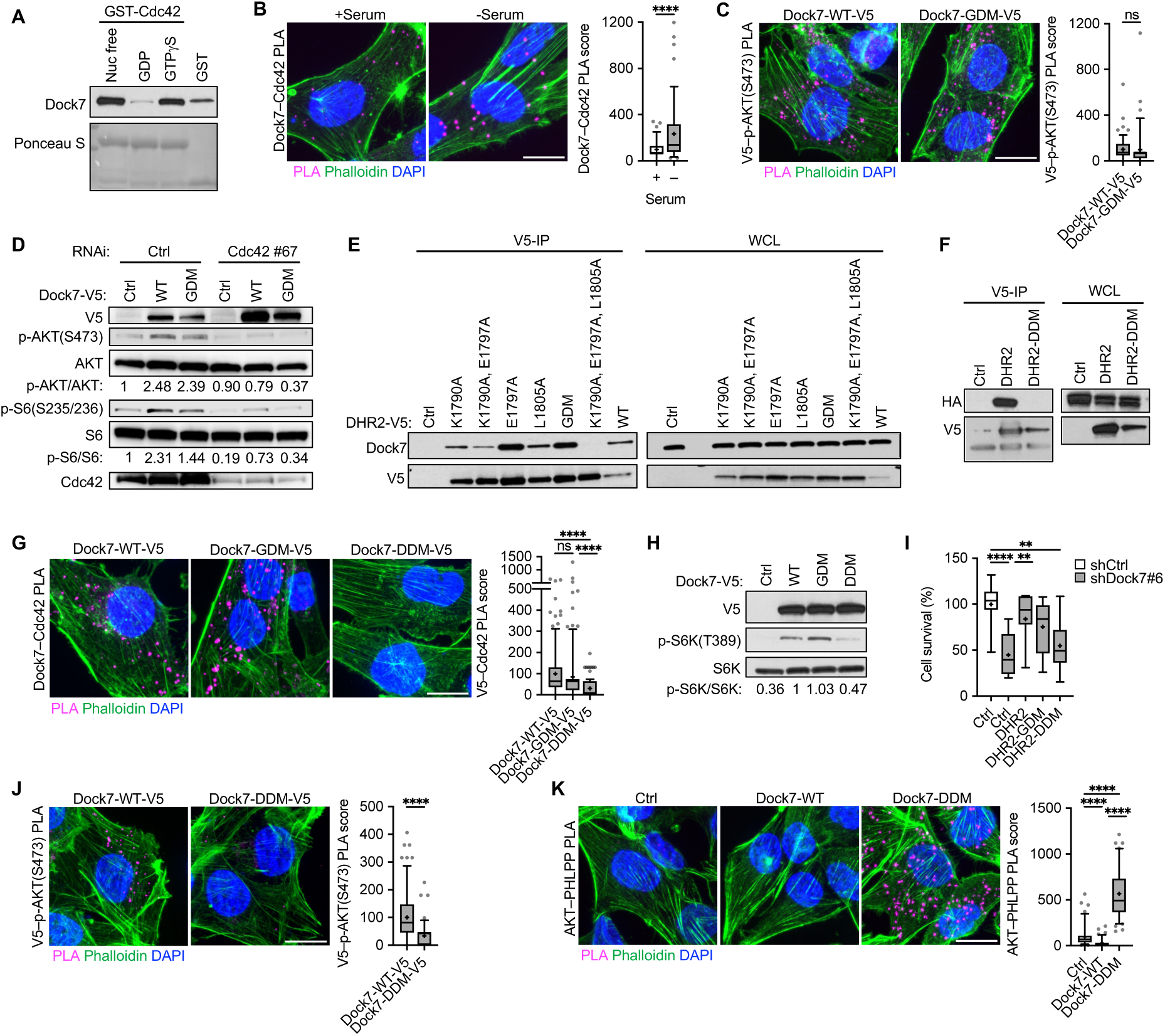
The role of the DHR2 domain in Dock7 function. (**A**) Recombinant GST-Cdc42 protein loaded with either GTPγS, GDP, or EDTA-treated were used to pull-down endogenous Dock7 from HEK-293T cells. Ponceau S was used to visualize the GST proteins. (**B**) PLA image and score of Dock7–Cdc42 interactions in MDA-MB-231 cells in full-serum or following overnight serum starvation. (**C**) PLA image and score of V5–p-AKT(S473) interactions in serum-starved MDA-MB-231 cells transiently overexpressing Dock7-WT-V5 and Dock7-GDM-V5. (**D**) Western blot of p-AKT and p-S6K from MDA-MB-231 cells transiently overexpressing Dock7-WT or Dock7-GDM while Cdc42 was knocked down using siRNA. (**E**) Generation of a Dock7 DHR2 dimerization domain mutant, DHR2-DDM where mutants semi-stably overexpressed in HEK-293T cells and immunoprecipitation studies were performed to isolate complexes between DHR2 and endogenous Dock7. (**F**) DHR2 was tagged with either HA or V5 and co-expressed in HEK-293T cells. Anti-V5 beads were used to immunoprecipitate V5-tagged DHR2 protein, and Western blots of HA to confirm interactions. (**G**) PLA image and score of V5–Cdc42 interactions in serum-starved MDA-MB-231 cells transiently overexpressing Dock7-WT-V5, Dock7-GDM-V5, and Dock7-DDM-V5. (**H**) Western blot of S6K activity in serum-starved HeLa cells either mock-transfected or transiently overexpressing either V5-tagged Dock7-WT, Dock7-GDM, or Dock7-DDM. (**I**) Cell survival of MDA-MB-231 cells following Dock7 knockdown and DHR2, DHR2-GDM, and DHR2-DDM overexpression after 2 days in serum-free media. (**J, K**) PLA images and score of (**J**) V5–p-AKT(S473) and **(K**) AKT–PHLPP interactions in serum-starved MDA-MB-231 cells transiently overexpressing Dock7-WT-V5 and Dock7-DDM-V5. Data shown as median ± interquartile range (box), 5^th^–95^th^ percentiles (whiskers), and mean (+).**P<0.01, ****P<0.0001, ns = not significant. Scale bar = 15 μm.

Based on analogy with other Dock proteins, three amino acid residues which, when mutated to alanine (K1759A, E1766A, L1774A), were predicted to generate a dimerization-defective mutant (DDM). We generated a series of DHR2 constructs with different combinations of point mutations and first tested the efficacy of these mutations to render DHR2 dimerization-defective by examining their ability to co-IP endogenous Dock7 in HEK-293T cells (Figure 6E). We identified the triple mutant as dimerization-defective and verified that dimer formation was occurring between two DHR2 domains by co-expressing a V5-tagged WT DHR2 domain with either WT DHR2-HA or DHR2-DDM-HA, and then immunoprecipitating the DHR2-V5. While DHR2-V5 was able to co-IP DHR2-HA, DHR2-DDM-HA was unable to do so, further demonstrating that the mutations disrupted dimerization within DHR2 (Figure 6F). Using PLA, we next examined the ability of Dock7-DDM-V5 to associate with Cdc42 in cells, as compared to Dock7-WT-V5 or Dock7-GDM-V5, in the absence of serum and found that the ability of Dock7-DDM to interact with Cdc42 was diminished (Figure 6G). These results indicate that mutations in the dimerization domain, which lies within the Cdc42 allosteric binding region, disrupt Cdc42-DHR2 as well as DHR2-DHR2 interactions.

Because of the requirement for Cdc42 in Dock7 function, we suspected that the Dock7-DDM, which had weakened affinity for Cdc42, would be compromised in its ability to promote a stress signaling response. Indeed, Dock7-DDM was less effective than Dock7-WT or Dock7-GDM in activating S6K under serum-free conditions (Figure 6H). Moreover, while DHR2 and DHR2-GDM overexpression following Dock7 knockdown was able to rescue cell survival, DHR2-DDM was less effective (Figure 6I). In PLA experiments, Dock7-DDM-V5 showed decreased V5–p-AKT(S473) interactions upon serum withdrawal compared to WT Dock7 (Figure 6J), suggesting the DHR2 dimer and/or Cdc42 is important for Dock7 interactions with p-AKT. PLA further demonstrated that the expression of Dock7-DDM increased interactions between AKT and PHLPP relative to WT Dock7 during serum starvation (Figure 6K).

### DockTOR complex assembly

PLA analyses suggest that two of the functionally important binding partners of Dock7, Cdc42 and p-AKT, have an increased capability for associating with Dock7 in cancer cells upon the serum withdrawal (Figures 6A and 3C, respectively), while pan-AKT–Dock7 interactions do not appear to be affected by the presence or absence of serum (Figure 3B). We further examined the relationships between the other Dock7-associated components and Dock7 using PLA in MDA-MB-231 cells. We found that endogenous Dock7 maintained interactions with mTOR in both serum-treated and serum-depleted cells, whereas Dock7–Rictor PLA signals decreased during serum starvation, suggesting a transient interaction between activated mTORC2 and the Cdc42-Dock7-mTOR complex (Figures 7A and 7B). The Dock7– TSC2 PLA signal was also similar in both serum and serum-free conditions suggesting that Dock7 interactions with TSC2 were serum-independent. However, Dock7–TSC1 interactions decreased upon serum depletion (Figures 7C and 7D), consistent with reports that TSC1 can dissociate from the TSC1/2 complex following an inactivating phosphorylation on TSC2 by AKT.^63–66^ Importantly, Rheb showed a significantly enhanced interaction with Dock7 when cells were deprived of serum (Figure 7E).

**Figure 7.**
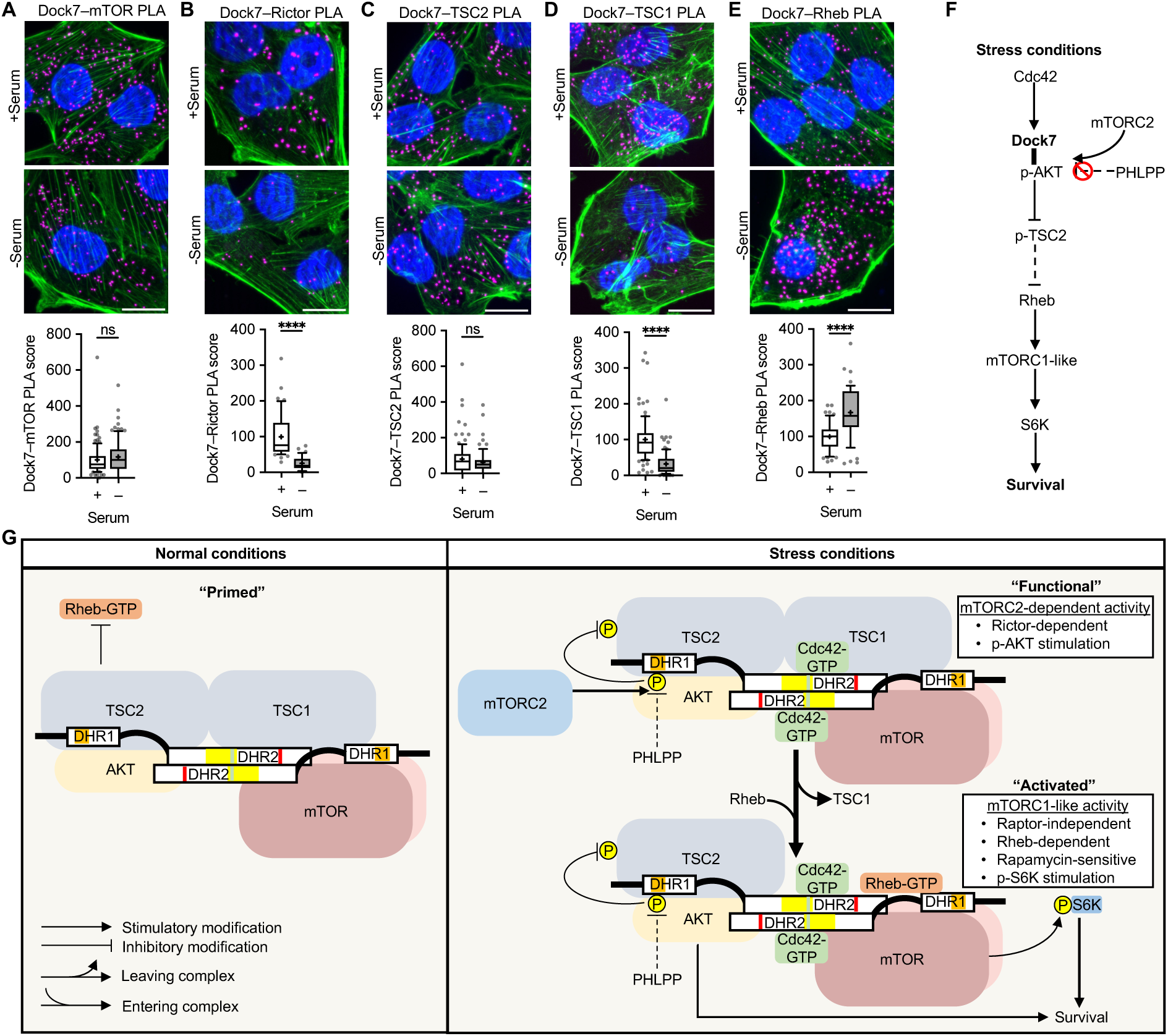
DockTOR complex assembly in response to serum deprivation. (**A**-**F**) PLA image and score for interactions between endogenous Dock7 and (**A**) mTOR, (**B**) Rictor, (**C**) TSC2, (**D**) TSC1, and (**E**) Rheb in MDA-MB-231 cells grown in complete media or serum-starved overnight. (**F**) Proposed model of Dock7-mediated signaling during stress conditions. (**G**) The DockTOR signaling complex during cellular response to stress. In complete media, the DockTOR complex is formed and “Primed”, and after the stress of serum withdrawal the complex is “activated” by mTORC2. Following mTORC2 activation of AKT, AKT phosphorylates TSC2 and TSC1 dissociates, disabling the Rheb GAP activity. In the “functional” DockTOR complex Rheb activates mTOR resulting in mTORC1-like activity and S6K activation. Data shown as median ± interquartile range (box), 5^th^–95^th^ percentiles (whiskers), and mean (+), or mean ± s.d. ****P<0.0001, ns = not significant. Scale bar = 15 μm.

Collectively, based on our data, we can define a series of signaling interactions (Figure 7F) that lead to a working model for the assembly and functioning of the Cdc42-Dock7-mTOR (Figure 7G). We propose a “core” multi-protein complex (DockTOR) that is promoted by GTP-bound Cdc42 binding to an allosteric regulatory site within Dock7, thus enabling Dock7 to function as a signaling scaffold that interacts with AKT, mTOR, TSC1, and TSC2 (Figure 7G, “primed”). Stress signals triggered by nutrient deprivation then begin a cascade of events which lead to activated mTORC2 catalyzing the phosphorylation of AKT at S473, followed by the phosphorylation of TSC2 at T1462 by AKT. Upon transitioning from a “functional” to an “activated” state, TSC1 dissociates from the complex disabling the GAP activity of TSC1/2 (Figure 7G). This transition enables Rheb-GTP to bind and stimulate mTOR within the DockTOR complex to mediate cell survival.

## Discussion

As our understanding of mTOR signaling has grown over recent years, new and alternative mTOR complexes in addition to mTORC1 and mTORC2 have been identified.^67–69^ Here, we describe an unconventional mTOR complex that plays an important role in the ability of cancer cells to survive when encountering stressful conditions including serum deprivation and the loss of a substratum. DockTOR acts as a pro-survival signaling node through its ability to maintain AKT phosphorylation under adverse conditions, ultimately propagating survival signals through AKT effectors including DockTOR-associated, Rapamycin-sensitive and Rheb-dependent mTORC1-like activity. Both Dock7 knockdown and AKT inhibition completely prevented cancer cells from exhibiting anchorage-independent growth and survival during serum deprivation. In mice, we find that Dock7 is critical for tumor formation, suggesting Dock7 may play an important role in supporting cancer survival during the more stressful stages of tumor progression such as tumor initiation from solitary tumor cells and seeding of secondary metastatic sites. Such tumor-initiating cells display a capacity to overcome various forms of stress and challenges such as hypoxia, nutrient deprivation, and oxidative stress that eliminate vulnerable populations of tumor cells.^70, 71^

We find that DockTOR stimulates mTOR/S6K in a Raptor-independent manner and does not appear to function at the lysosome. Raptor-independent S6K recruitment can occur through interactions with the FKBP-Rapamycin binding (FRB) domain of mTOR that place S6K near the active kinase domain.^72^ This allows for Rapamycin-sensitivity to be maintained in the absence of Raptor, as Rapamycin forms an inhibitory complex with FKBP12 and subsequently binds to the FRB domain of mTOR.^73^ The mTORC2 component, Rictor, partially obscures the FRB domain of mTOR, which gives rise to the insensitivity of mTORC2 activity to Rapamycin.^74^ Therefore, our finding that Dock7 stimulation of mTOR/S6K during stress conditions requires Rictor and is Rapamycin-sensitive indicates that there are both Rictor-dependent (mTORC2) and Rictor-independent (mTORC1-like) activities necessary for DockTOR function, thus providing for a distinctive mTOR signaling switch. While there have been reports of Rapamycin-sensitive but Raptor-independent mTORC1-like activities,^75^ what makes DockTOR unique is that by lacking Raptor it allows mTOR/S6K activation during nutrient-deprived conditions when canonical mTORC1 is inhibited by the AMPK-dependent phosphorylation of Raptor^76^. Hyperactivated mTORC1 has been shown to inhibit apoptosis during serum deprivation^77^ and the sustained mTORC1-like activity maintained by DockTOR provides an important survival benefit during such stress conditions. However, S6K knockdown reduced the ability of cancer cells to grow in soft agar by half, indicating S6K activity is an important, but not an exclusive AKT signaling output in the DockTOR pro-survival response, as AKT triggers additional downstream signals that block apoptosis.

DockTOR incorporates a previously undefined role for the DHR1 domain in the ability of Dock7 to preserve the mTORC2-catalyzed phosphorylation of AKT at Ser473. While the low-level of AKT activity that is maintained by the DHR1 domain upon the removal of serum from culture conditions may have previously been considered “background,” it is in fact critically important for cancer cells to resist anoikis and apoptosis induced by nutrient deprivation. The protection afforded by Dock7 to the phosphorylation of AKT at Ser473 appears to occur through an unconventional mechanism specific to the C2-like motif of its DHR1 domain, as C2 and C2-like motifs have not previously been implicated in binding to phospho-proteins. We show that the C2-like motif interacts with phosphorylated AKT and protects it from dephosphorylation as underscored by the observation that loss of a functional C2-like motif enhances interactions between AKT and its phosphatase PHLPP. Previously, very little had been known about the DHR1 domain of the Dock7, so we were surprised to find that it rescued anchorage-independent growth, cell survival, and signaling to S6K in cells depleted of Dock7. The ability of the DHR1 domain to promote AKT activity by preserving its mTORC2-catalyzed phosphorylation state may promote a sufficient activation of mTOR and/or trigger other AKT-dependent survival signals to restore transformed phenotypes that are lost when depleting cells of endogenous Dock7.

We have found that a DHR1 construct containing a C-terminal 183 amino acid extension (DHR1L) is more effective than the exact limit DHR1 domain in restoring the signaling functions lost when Dock7 is knocked down, suggesting that this linker region extension may play an important role in binding to the other signaling partners comprising DockTOR. Given that DHR1L partially rather than fully restores the actions of Dock7 in transformation assays and xenograft experiments, underscores the importance of additional regions within the full-length Dock7 protein, including DHR2, to generate a complete signaling-competent complex. Interestingly, the DHR2 domain also provides some compensation for the loss of Dock7, which is not due to its GEF domain as the GEF-defective mutant is as effective in these assays as the wild-type protein, whereas a dimerization-defective DHR2 mutant is ineffective. While DHR2 showed lower interactions with p-AKT(S473) compared to DHR1L, the expression of DHR2 still helped to prevent interactions between AKT and PHLPP. It may be that the ability of the DHR2 dimer to bind AKT sterically blocks the access of phosphatases to maintain a degree of AKT activation.

Several intriguing questions for future study are raised by this first report of a stress-dependent Cdc42-Dock7-mTOR signaling complex. One involves the mechanism responsible for the initial activation of Cdc42. Dock7 has been mainly studied in brain development where it regulates neuronal polarity and Schwann cell migration^78, 79^ and initial studies suggested a similar classical role for Dock7 as a GEF for Cdc42/Rac in cancer.^80–82^ However, we find that the Dock7 GEF activity is dispensable for the Dock7-mediated activation of AKT and mTOR/S6K. GEF-independent Dock7 functionality has been previously described^83, 84^ and may stem from the fact that Dock7 serves a more impactful role as a scaffolding protein in this context. It remains to be seen if other GEFs are involved in promoting the activation of Cdc42 and/or if sufficient pools of GTP-bound Cdc42 are generated prior to serum deprivation or other stresses that enable Cdc42 to bind to the allosteric site within the DHR2 domain of Dock7 where its activated state can be stabilized, as occurs when Cdc42 binds its target/effector proteins.^85^ Another interesting question concerns where DockTOR assembles in cells, especially given that Raptor is not part of the complex and because we find that DockTOR is not localized to lysosomes.

Ultimately, it will be important to elucidate the structural features of the different protein-protein interactions comprising the DockTOR complex as well as determine the identity of its downstream signals through proteomic and phospho-proteomic studies to define the global changes that are promoted by the DockTOR signaling axis during serum starvation to allow for cancer cell survival. Such studies will help to better distinguish the functions and molecular mechanisms of this distinct signaling node from the previously described growth factor-dependent AKT and mTOR pathways. Increasing evidence suggests that there may be additional mTOR-interacting proteins in different tissues and cell lines,^68, 86^ and it has become apparent that mTOR signaling is more complex than originally thought. Understanding how mTOR signaling utilized by cancer cell allows them to survive the challenges they face during tumor progression due to anoikis, nutrient deprivation, and other forms of stress could highlight previously unappreciated vulnerabilities and open the way toward designing new therapies.

## Acknowledgements

This research was supported by grants from the NIH (R35GM152206 and CA201402) to R.A.C., NSF Graduate Research Fellowship (DGE-1650411) to O.Y.T.P., and NCI/NIH Ruth L. Kirschstein NRSA under Award No. F32CA275151 to M.R.Z.

## Author contributions

O.Y.T.P., M.R.Z., M.J.L., K.F.W., and R.A.C. designed the experiments; O.Y.T.P., M.R.Z., M.J.L., R.R.S., K.S.R., and I.Y.H. performed experiments; O.Y.T.P., M.R.Z., M.J.L., R.R.S, and I.Y.H. analyzed data; O.Y.T.P., M.R.Z., K.F.W., and R.A.C. wrote the manuscript; K.F.W. and R.A.C. supervised the project; and all authors revised the manuscript and approved the final version.

## Author declarations

The authors declare no competing interests.

## Materials and Methods

### Cell lines, cell culture, and reagents

HEK-293T and all cancer cells were obtained from American Type Cell Culture Collection (ATCC), except TSE breast cancer cells (kindly supplied by Dr. Steven Abcouwer, University of Michigan), and MDA-MB-231 breast cancer cells metastasized to brain (kindly supplied by Dr. Joan Massagué, MSKCC^87^). Cells were maintained at 37°C, 5% CO_2_ in DMEM (Thermo Fisher) supplemented with 10% fetal bovine serum (FBS; Thermo Fisher). All cell lines were tested and found negative for mycoplasma contamination. For growth factor stimulation, 7×10^5^ cells were seeded in 100 mm dishes (Corning), serum-starved for 18-24 h, then stimulated with Heregulin β (HRG), EGF domain, residues 178-241 (Sigma-Aldrich) at the concentration and times indicated, followed by cell lysis. For mTOR/AKT inhibitor analysis, 1×10^6^ cells were seeded in 100 mm dishes, allowed to recover overnight, and then treated with either vehicle DMSO, 1 nM Rapamycin (Cell Signaling Technologies, #9904), 100 nM Torin (Cell Signaling Technologies, #14379), or 10 μM MK2206 (SelleckChem, #S1078) in serum-free DMEM for 18-24 h. For phosphatase inhibitor analysis, 1×10^6^ cell were seeded in 100 mm dishes, allowed to recover overnight, and then serum-starved for 18-24 hours, cells were then treated with either DMSO, 10 nM Okadaic Acid (Cell Signaling Technologies, #5934), or 50 nM Calyculin A (Cell Signaling Technologies, #9902) in serum-free media for 1 h before collection and lysis. For soft agar drug treatment, the same concentration of inhibitor was used, but the DMEM media was supplemented with 10% FBS and 1x Pen/Strep.

### DNA constructs, siRNA, and shRNA

Rac1, Cdc42, Rheb, Dock7, wild-type and point mutation constructs used for transient transfections were cloned in our laboratory into pcDNA3.1 (Thermo Fisher). Full length Dock7 was cloned using the pcDNA™3.1 Directional TOPO™ Expression Kit (Invitrogen). RNA was isolated from Hela cells, cDNA synthesized with SSII Reverse Transcriptase and the mammalian expression plasmid was generated containing a C-terminal V5 epitope tag. Whole plasmid sequencing revealed that this full length Dock7 sequence was consistent with isoform 4 (NM_001272001.2), and this was used to create mutant and truncation constructs to complete this study. TSC1, TSC2, mTOR, and AKT DNA constructs were obtained from Addgene (plasmid #12133, #14129, #1861, #9021, respectively).^24, 63, 65^ Constructs used for transient transfections were cloned into pcDNA3.1 (Thermo Fisher, V79020) and site-directed mutagenesis was performed to create point mutation variant constructs using the QuikChange Site-Directed Mutagenesis Kit (Agilent Technologies, #200518). Lentiviral constructs of Dock7, Dock7 truncations, Cdc42, Rheb, and YFP used were cloned into pSIN-EF2. Constructs of Dock7 contained the following amino acids: DHR1E (561-727), DHR1L (561-910), and DHR2 (1540-2098). GST-Rheb and GST-Cdc42 was cloned into pGEX (GE Healthcare Life Sciences). Silencer Select siRNAs of Rheb (s12019, s12020, s12021) and Cdc42 (s2765, s2766, s2767) were purchased from Thermo Fisher. shRNA for Dock7 (shRNA1: TRCN0000377466, shRNA2: TRCN0000376575, shRNA3: TRCN0000365143, shRNA6: TRCN0000365145), Raptor (shRNA1: TRCN0000010415, shRNA2: TRCN0000010416), and Rictor (shRNA1: TRCN0000074288, shRNA2: TRCN0000074289) were purchased from Sigma.

### Transfection

5×10^5^ cells were seeded in 100 mm dishes (Corning) then transfected with 4 μg DNA using Lipofectamine and Plus Reagent (Thermo Fisher) according to the manufacturer’s protocol. Cells recovered in complete medium for 3 h followed by serum starvation for 18-24 h. For knockdown experiments, cells were seeded and transfected with 2.5 nM siRNA the next day using Lipofectamine2000 (Thermo Fisher) following the manufacturer’s protocol. For rescue experiments, after siRNA transfection cells were then split onto 60 mm dishes at 2.5×10^5^ cells/dish, allowed to recover overnight, and transfected with 1 μg of DNA using Lipofectamine and Plus Reagent. For HEK-293T cells, 3×10^6^ cells were seeded in 100 mm dishes, cultured for 24 h, and transfected with 4 μg DNA the next day using Lipofectamine and Plus Reagent following the manufacturer’s protocol.

### Lentiviral transduction

To generate lentivirus, HEK-293T cells were cultured for 24 h in 100 mm dishes to 80% confluency and transfected with constructs. 6 μg of construct DNA (pSIN for overexpression or pLKO for shRNA), 4 μg pCMV, and 2 μg pMD2.G were added to 800 μl serum-free DMEM and mixed with 30 μl PEI. DNA/PEI/DMEM solution was incubated for 15 min at room temperature, then added to cells in 12 ml of fresh media and cultured overnight. Media was then changed to 13 ml of complete media, cells were cultured for 24 h, and spent media was collected. To harvest virus, spent media was centrifuged at 1000 rpm for 10 min to remove cell debris, sterile filtered using a 0.45 μm syringe filter. 13 ml media was replaced and after 24 h virus was collected again. Collected virus was combined, mixed, pipetted into 3 ml aliquots, and stored at ࢤ80°C. For lentiviral transduction, 7×10^5^ target cells were seeded in 100 mm plates, cultured overnight, and lentivirus with polybrene (1:1000) in complete media was added. Cells were cultured overnight, washed with 1x PBS, cultured for 48 h in complete media, then passaged. Cells expressing constructs were selected with 2 μg/ml puromycin for 3-5 days to create semi-stable lines and maintained in complete media supplemented with 1 μg/ml puromycin.

### CRISPR-Cas9 knock-out

Dock7 was genetically ablated using CRISPR-Cas9 knock-out plasmid and HDR plasmid transfection. Cells were seeded in 6-well plates at 2.5×10^5^ cells/well in antibiotic-free complete media, allowed to recover overnight, then treated. Solutions of 1 μg CRISPR (Santa Cruz Biotechnology, sc-404461) and DOCK7 HDR (h) (Santa Cruz Biotechnology, sc-404461-HDR) plasmid DNA were prepared and mixed, then added to 150 μl Plasmid Transfection Medium (Santa Cruz Biotechnology, sc-108062). A separate solution of 5 μl UltraCruz Transfection Reagent (Santa Cruz Biotechnology, sc-395739) in 150 μl Plasmid Transfection Medium was also prepared and mixed, then both solutions were incubated for 5 min. Solutions were mixed, incubated for 15 min, and added to cells. Media was changed after 24 h, then after 48 h selection was performed with 1.75 μg/ml puromycin.

### Immunoblot analysis

Cells were lysed with cell lysis buffer (50 mM Hepes pH 8.0, 150 mM NaCl, 1 mM MgCl_2_, 25 mM NaF, 1 mM Na_3_VO_4_, 50 mM β-glycerophosphate, 10 μg/ml Leupeptin, 10 μg/ml Aprotinin, and 1% Triton X-100). The lysates were resolved by SDS-PAGE (4-20% Tris-Glycine gels, Thermo Fisher), and then the proteins were transferred to polyvinylidene fluoride (PVDF) membranes (PerkinElmer). The membranes were incubated with the indicated primary antibodies diluted In 20 mM Tris pH 7.4, 150 mM NaCl, and 0.05% Tween-20. Primary antibodies were detected with horseradish peroxidase-conjugated secondary antibodies (Cell Signaling Technology) followed by exposure to ECL reagent (PerkinElmer). Dock7 and Flag antibodies were purchased from Sigma. pan-mTOR, pan-S6 kinase, Cool-1, Rac1, and Cdc42 antibodies were obtained from Millipore. Myc and HA antibodies were from Covance and anti-actin was purchased from NeoMarker. Other antibodies were purchased from Cell Signaling Technology.

### Immunoprecipitation

HEK-293T cells were lysed with lysis buffer (50 mM Hepes pH 8.0, 150 mM NaCl, 1 mM MgCl_2_, 25 mM NaF, 1 mM Na_3_VO_4_, 50 mM β-glycerophosphate, 10 μg/ml Leupeptin, 10 μg/ml Aprotinin, 0.3% CHAPS). Lysates were pre-cleared with BSA-coated Protein G beads (Thermo Fisher) on a rotator at 4°C for 15 min. The supernatant was collected and added with anti-Myc (Covance), HA (Covance), Flag (Sigma-Aldrich), or V5 (Sigma-Aldrich) antibody for 2 h, then BSA-coated Protein G beads were added and incubated for 1 h at 4°C. Immunoprecipitates were washed 3x with lysis buffer, 2x Laemmli sample buffer was added, and analyzed on 4-20% Tris-Glycine gels followed by immunoblot analysis as described above.

### Blue Native PAGE

Blue Native PAGE was performed using lysates prepared from growing HEK-293T cells according to the manufacturer’s protocol (Thermo Fisher). 1×10^7^ cells were collected and lysed using 1x Native PAGE lysis buffer including protease inhibitor cocktail and 1% digitonin. Multiple lanes of the same lysates were then run on one 3-12% Bis-Tris gel followed by denaturing treatment and transferred onto PVDF membrane. Each individual strips of lysates were then blotted for mTOR, Dock7, TSC1, and TSC2. All antibodies were purchased from Cell Signaling Technology. Blots were developed as described above followed by processing and compilation using ImageJ.^85^

### Nucleotide-dependent GST fusion protein pull-down

GST, GST-Rheb, and GST-Cdc42 were expressed in BL21 cells and purified by affinity chromatography using Glutathione Sepharose High Performance beads (GE Healthcare Life Sciences, #17527901) according to the manufacturer’s protocol. Purified protein was stored on the glutathione beads with 30% glycerol at ࢤ20°C until use. HEK-293T whole cell lysates were lysed with cell lysis buffer (50 mM Hepes pH 8.0, 150 mM NaCl, 1 mM MgCl_2_, 25 mM NaF, 1 mM Na_3_VO_4_, 50 mM β-glycerophosphate, 1 mM DTT, 10 μg/ml Leupeptin, 10 μg/ml Aprotinin, 1% Triton X-100). Lysates were pre-cleared using GST beads. GST-Rheb and GST-Cdc42 was treated with 10 mM EDTA, 10 mM EDTA + 1 mM GDP, or 10 mM EDTA + 100 μM GTPγS at room temperature for 15 min followed by the addition of 50 mM MgCl_2_ for the samples containing nucleotides. GST controls were either treated with EDTA or non-nucleotide loaded (no EDTA). Nucleotide-loaded or nucleotide-free GST/GST-Rheb/GST-Cdc42 were added to the pre-cleared lysates. Pull-downs were done in the lysis buffer ± EDTA at 4 °C for 2 h. Pulled-down proteins were washed 3x with the lysis buffer ± EDTA, 2x Laemmli sample buffer was added, and analyzed on 12% Tris-Glycine gels followed by immunoblot analysis as described above.

### Soft agar colony formation

HeLa, SK-BR-3, MCF7, and MDA-MB-231 cells in complete media containing 0.3% agarose were seeded at 0.8-1×10^4^, 5×10^3^, 1×10^4^ and 2×10^4^ cells/well, respectively, onto a layer of 0.6% agarose with complete media in 6-well plates. Cultures were fed every 3-4 days with complete medium containing 0.3% agar. After 14-21 days, 1 mg/ml NBT in 1x PBS was added, cultured overnight, and colonies were counted.

### Cell Counting Kit (CCK)-8 assay

Cells were seeded in 96-well plates at 16,000 cells/well and cultured overnight in 100 μl of complete media. The next day, cells were washed with 1x PBS and media was changed to serum-free media, and media was replaced every day. The CCK8 assay (Dojindo Laboratories, CK04) was then performed each day according to the manufacturer’s instructions. Briefly, 10 μl of CCK8 solution to each well, plates were incubated for 4 h, and the absorbance of each well was measured at 450 nm using a Tecan SPARK microplate reader.

### Cell survival assay

Cells were seeded in 48-well plates at 16,000 cells/well and cultured overnight in complete media. The next day, cells were washed with 1x PBS and media was changed to serum-free media, with fresh media replaced every day. After two days cells were then trypsinized and counted. Counts were normalized to control conditions.

### EdU proliferation assay

DNA synthesis was directly measured in live cells with the EdU Staining Proliferation kit iFluor 488 (Abcam, ab219801) following the manufacturer’s protocol. Cells were seeded in 6-well plates with no. 1.5 22×22 mm coverslips (Electron Microscopy Sciences, 72204-01) at 16,000 cells/well, cultured for 48 h, then washed with 1x PBS and treated with complete or serum-free media. After a 18-24 incubation, cells were incubated with 10 μM EdU solution for 4 h, fixed in 4% paraformaldehyde (PFA; Thermo Fisher, J19943.K2) for 10 min, permeabilized, and labeled using kit components. Coverslips were mounted using Vectashield with DAPI (Vector, H-1200) on microscope slides and sealed. The ImageJ Particle Analyzer plugin was used to calculate proliferation as the fraction of EdU positive cells.

### Cancer cell spheroid invasion

MDA-MB-231 cancer cell spheroids were generated as previous described.^88^ Spheroids were encapsulated in 2.5 mg/ml collagen (Corning) and then spheroid invasion was monitored over 24-48 h post-embedding.

### TUNEL assay

Cells were seeded in 6-well plates with no. 1.5 22×22 mm coverslips at 16,000 cells/well, cultured for 48 h in complete media, then washed with 1x PBS, and then treated with complete or serum-free media for 18-24 h. After the incubation, media was removed and cells were washed 3x with 1x PBS, fixed in 4% PFA (Thermo Fisher, J19943.K2) for 10 min, washed 3x with 1x PBS, and stored in 1x PBS at 4°C until the In Situ Cell Death Detection Kit, Fluorescence (Roche, 11684795910) was used according to the manufacturer’s protocol. Analysis was performed in ImageJ using the Particle Analyzer plugin.

### Immunofluorescence staining

Cells were seeded in 6-well plates with no. 1.5 22×22 mm coverslips at 16,000 cells/well, cultured for 48 h, washed with 1x PBS, and treated with complete or serum-free media for 18-24 h. Samples were then washed with 1x PBS, fixed in 4% PFA for 10 min, and washed 3x with 1x PBS. Samples were permeabilized with a 15 min wash in 0.1% Triton X-100, washed 3x with 0.02% Tween20/1x PBS for 15 min each, then blocked in 0.02% Tween20/3% BSA/10% FBS for 4 h. After blocking, samples were incubated in primary antibodies for mouse anti-Dock7 (1:200; Santa Cruz Biotechnology, sc-398888) and rabbit anti-LAMP1 (1:200, Cell Signaling Technology, 9091S) overnight at 4°C. Samples were next washed 3x with 0.02% Tween20/1x PBS for 15 min each and incubated with goat anti-mouse AlexaFluor 488 (Invitrogen, A11029) and goat anti-rabbit AlexaFluor 594 (Invitrogen, A11037) secondary antibodies for 2 h at room temperature. Following 3 washes with 0.02% Tween20/1x PBS for 15 min each, and coverslips were mounted on a microscope slide using Vectashield mounting media with DAPI and sealed.

### Proximity ligation assay (PLA)

Cells were seeded in 6-well plates with no. 1.5 22×22 mm coverslips at 16,000 cells/well, cultured for 48 h, washed with 1x PBS, and treated with complete or serum-free media for 18-24 h. After incubation, cells were washed 3x with 1x PBS, fixed in 4% PFA (Thermo Fisher, J19943.K2) for 10 min, washed 3x with 1x PBS, and stored in 1x PBS at 4°C until use. Samples were permeabilized with 0.1% Triton X-100 (Millipore, 1.08603.1000) for 15 min, washed 3x for 15 min with 0.02% Tween20 (Sigma-Aldrich, P1379) in 1x PBS, then blocked in Duolink Blocking Solution (Sigma-Aldrich, DUO82007) for 60 min at 37°C. Primary antibodies were diluted in Duolink Antibody Diluent (Sigma-Aldrich, DUO82008) and incubated overnight at 4°C (Supplementary Table S1). PLA was next performed using Duolink In Situ PLA Probe Anti-Mouse PLUS (Sigma-Aldrich, DUO92001), Duolink In Situ PLA Probe Anti-Rabbit Minus (Sigma-Aldrich, DUO92005), and In Situ Detection Reagents Red (Sigma-Aldrich, DUO92008) following the Duolink PLA Fluorescence protocol. In Situ Wash Buffers Fluorescence (Sigma-Aldrich, DUO82049) were used for all wash steps in PLA procedures. Following PLA, cells were counterstained with AlexaFluor 488 Phalloidin (1:200; Invitrogen, A12379) overnight at 4°C. Coverslips were then mounted on a microscope slide using Vectashield mounting media with DAPI and sealed. When examining LAMP1 immunofluorescence with Dock7–mTOR PLA, PLA was performed followed by immunofluorescence as described above. Analysis was performed in ImageJ, and PLA puncta were counted using the Find Maxima plugin. PLA scores were determined by normalizing the PLA spot count per cell to the average number of PLA spots in control cells, which was set to 100.

### Phase contrast microscopy

To evaluate spheroid outgrowth, phase contrast imaging was performed using using a Keyence BZ-X810 inverted fluorescence phase contrast microscope using a Plan Apochromat 10x/0.45 NA objective (BZ-PA60, Keyence Corp).

### Fluorescence microscopy

maging was performed on a Keyence BZ-X810 inverted fluorescence phase contrast microscope (BZ-PA60, Keyence Corp) and ET DAPI (ex. 395/25 em. 460/50; Chroma, 4900-UFI), ET EGFP (ex. 470/40 em. 525/50; Chroma, 49002-UFI), and ET mCH/TR (ex. 560/40 em. 630/75; Chroma, 49008-UFI) filters was used for fluorescence imaging. PLA images were taken with a Plan Apochromat 60x/1.4 NA oil immersion objective using optical sectioning (1D, width=10) and z-stacks (0.5 μm pitch) taken through the height of the cells. EdU and TUNEL images were taken using a 20x/0.75 NA objective.

### Isolation of Mouse Embryo Fibroblasts (MEF)

Dock7 heterozygous mice were crossed, and embryos were isolated at dpc13.5 to create Mouse Embryonic Fibroblasts (MEFs). Pregnant mice were euthanized as approved by AICUC 14 days after appearance of copulation plug. The uterus was removed, and embryos were extracted and placed in 60 mm tissue culture dishes. The head was removed and used for genotyping, and internal organs were discarded. Each embryo was washed with HBSS, without calcium nor magnesium (Thermo Fisher, #14170120) twice, and then placed in 0.25% trypsin-EDTA (Corning, #25053CI). Embryonic tissue was minced into small pieces, and incubated at 5% CO_2_, 37°C for 5 min. The mixture was pipetted up and down and placed back in the incubator for 10 min, trypsin was then deactivated with 15 mL DMEM supplemented with 10% FBS and transferred to a 50 ml tube where it rested for 10 min. Single cells and the cell cluster supernatant were transferred to a 100 mm plate, cells attached overnight, and media was changed to DMEM with 10% FBS after 24 h. Cells were used 48 h later.

### Tumor xenograft

The different conditions of MDA-MB-231 breast cancer cells were pre-prepared and 3×10^6^ cells were injected subcutaneously into each flank of female NSG mice, aged 8 weeks. Tumor size was measured using calipers every 7 days, and tumor volume was calculated using the formula 0.5 x (length x width^2^). Tumors from NSG mice were harvested 56 days after the xenograft inoculation. All mice experiments were carried out according to the protocols (#2003-0097) approved by the Center for Animal Resources and Education (CARE) at Cornell University. Tumor formation was assessed as the time when tumor reached 10 mm^3^.

### TCGA Data

The Cancer Genome Atlas (TCGA) Breast Cancer (BRCA) data set was assessed and analyzed through UCSC Xena browser 87.

### Statistical analysis

Statistical analysis was performed using GraphPad Prism 9.0. Normality in was tested with the D’Agostino-Pearson omnibus normality test. Data with a Gaussian distribution were compared using a two-tailed Student’s t-test for two groups and one-way ANOVA with Dunn’s post-hoc analysis for multiple groups. Non-parametric tests were used to compare data with a non-normal distribution and a two-tailed Mann-Whitney test was used for two groups, while one-way Kruskal-Wallis test was used for multiple groups. No statistical method was used to determine sample size. All experiments were reproduced at least three independent times.

## Notes

### Competing Interest Statement

The authors have declared no competing interest.

### Summary of Updates

Additional data added to the manuscript and supplemental files

