## Supplementary Figures and Tables for "A multiprotein signaling complex sustains AKT and mTOR/S6K activity necessary for the survival of cancer cells undergoing stress"

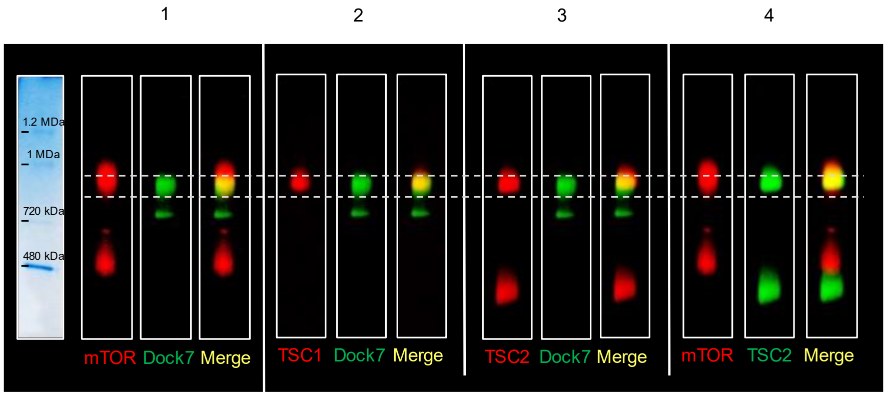


- - - Marks the native molecular weight of the majority of Dock7 species

Co-migration (same native molecular mass) of the two proteins shown within a panel

**Figure S1. Dock7 co-migrates with mTOR and its main negative regulator, the TSC1/2 complex.** HEK-293 cells grown in complete medium were lysed and native complexes were separated using Blue Native-PAGE (BN-PAGE). Each lane was stained by the specified antibodies.

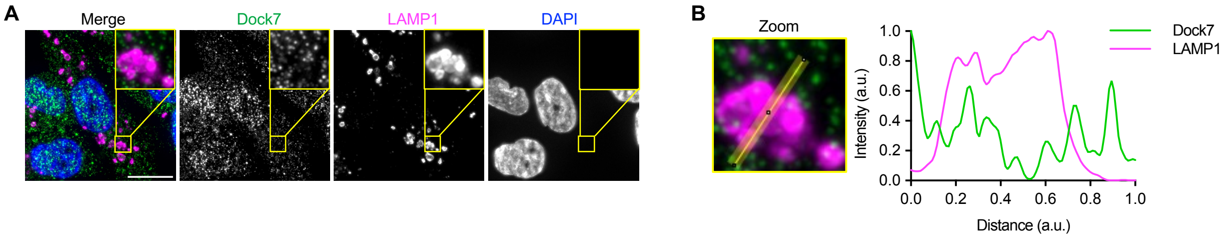


**Figure S2. Dock7 does not colocalize with lysosomal-marker LAMP1.** (**A**) Immunofluorescence images of Dock7 and LAMP1 in MDA-MB-231 cells in serum-free media. Insert show boxed region of interest (ROI). (**B**) Zoomed ROI and line-scan of Dock7 and LAMP1 fluorescence intensity. Scale bar = 15 µm.

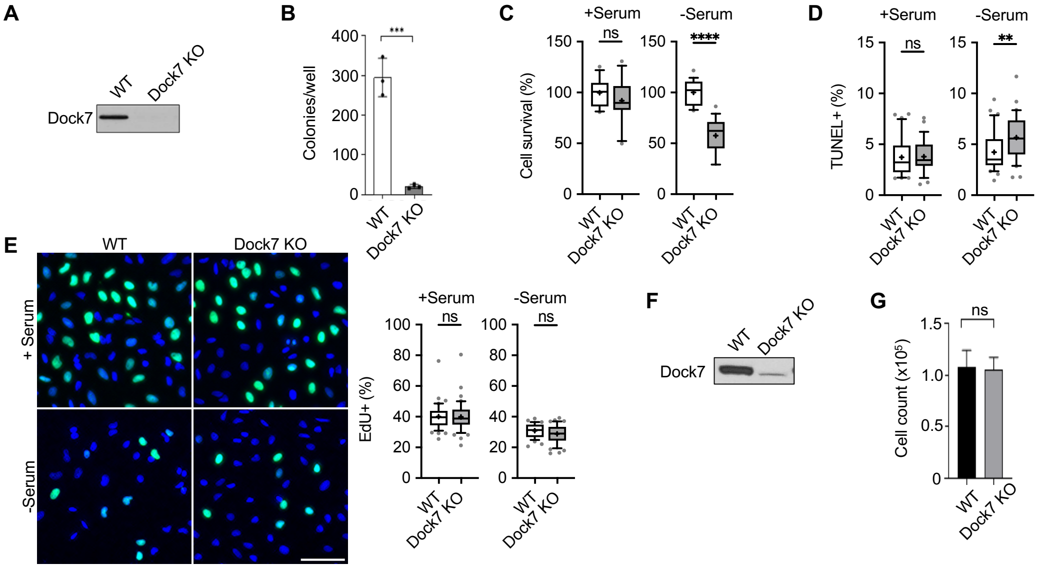


**Figure S3. HeLa Dock7 knock-out cell line.** (**A**) Western blot of Dock7 expression in HeLa WT and Dock7 KO cell lines. (**B**) Soft agar colony formation of HeLa WT and Dock7 KO cells. (**C**) Cell survival of HeLa WT and Dock7 KO cells in serum-free media after 2 days. (**D**) TUNEL assay of HeLa WT and Dock7 KO cells cultured in complete media or serum-starved overnight. (**E**) EdU assay to determine cell proliferation of HeLa WT and Dock7 KO cells performed in complete media or after overnight serum starvation. (**F**)Western blot of Dock7 protein expression of Dock7 KO MEF cell line. (**G**) Cell survival of WT and Dock7 KO MEF cells cultured in serum-free media for 2 days. Data shown as median ± interquartile range (box), 5$^{th}$–95$^{th}$ percentiles (whiskers), and mean (+), or mean ± s.d. **P<0.01, ****P<0.0001, ns = not significant. Scale bar = 100 μm.

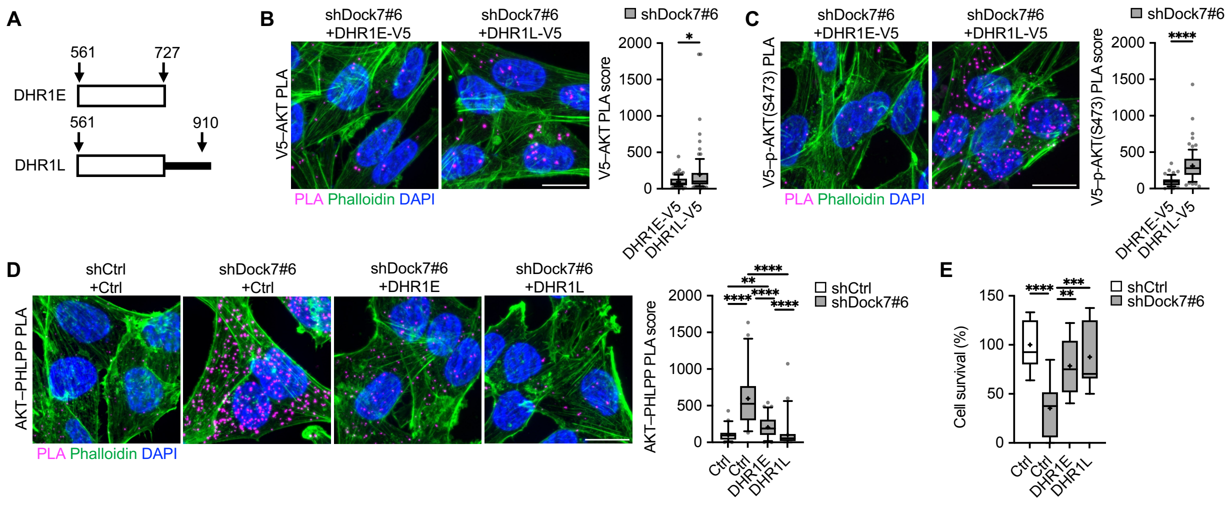


**Figure S4. DHR1L is more effective than DHR1E at rescuing function following Dock7 knockdown.** (**A**) Schematic of DHR1E and DHR1L constructs. (**B**) Cell survival of MDA-MB-231 cells following Dock7 knockdown and DHR1E and DHR1L overexpression after 2 days in serum-free media. (**C**-**E**) PLA image and score of (**C**) V5–AKT, (**D**) V5–p-AKT(S473), and (**E**) AKT–PHLPP interactions in MDA-MB-231 cells after overnight serum starvation following Dock7 knockdown and DHR1E and DHR1L overexpression. Data shown as median ± interquartile range (box), 5$^{th}$–95$^{th}$ percentiles (whiskers), and mean (+). *P<0.05, **P<0.01, ***P<0.001 ****P<0.0001, ns = not significant. Scale bar = 15 μm.

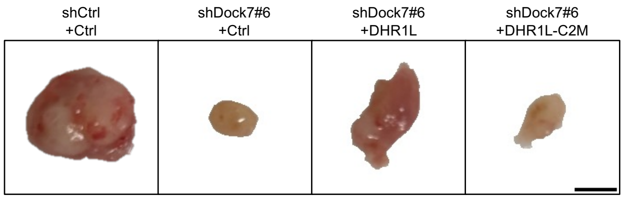


**Figure S5. NSG xenograft tumor growth after 8 weeks.** Representative images of NSG mouse tumor xenograft of MDA-MB-231 cells following Dock7 knockdown and DHR1L and DHR1L-C2M limit domain overexpression after 8 weeks. Scale bar = 5 mm.

**Table S1. PLA antibodies.**

| Pair | Ms antibody | Rb antibody |
|---|---|---|
| Dock7–TSC1 | Dock7 (1:100; Santa Cruz Biotechnology, sc-398888) | TSC1 (1:100, Cell Signaling Technology, 6935S) |
| Dock7–TSC2 | Dock7 (1:100; Santa Cruz Biotechnology, sc-398888) | TSC2 (1:100; Cell Signaling Technology, 4308S) |
| Dock7–mTOR | Dock7 (1:100; Santa Cruz Biotechnology, sc-398888) | mTOR (1:100; Cell Signaling Technology, 2983S) |
| Dock7–Rheb | Dock7 (1:100; Santa Cruz Biotechnology, sc-398888) | Rheb (1:100, Cell Signaling Technology, 13879S) |
| Dock7–Raptor | Dock7 (1:100; Santa Cruz Biotechnology, sc-398888) | Raptor (1:100; Cell Signaling Technology, 2290S) |
| Dock7–Rictor | Dock7 (1:100; Santa Cruz Biotechnology, sc-398888) | Rictor (1:100; Cell Signaling Technology, 2114S), |
| Dock7–LAMP1 | Dock7 (1:100; Santa Cruz Biotechnology, sc-398888) | LAMP1 (1:100, Cell Signaling Technology, 9091S) |
| Dock7–AKT | Dock7 (1:100; Santa Cruz Biotechnology, sc-398888) | AKT (1:100; Cell Signaling Technology, 9272S) |
| AKT–PHLPP | AKT (1:100, Cell signaling Technology, 2920S) | PHLPP (1:50, Proteintech, 22789-1-AP) |
| Dock7–p-AKT(S473) | p-AKT(S473) (1:50, Cell Signaling Technology, 4051S) | Dock7 (1:100; Abcam, ab118790) |
| p-AKT(S473)–p-TSC2(T1462) | p-AKT(S473) (1:50, Cell Signaling Technology, 4051S) | p-TSC2(T1462) (1:50, Cell Signaling Technology, 3617S) |
| V5–AKT | V5-tag (1:100, Cell Signaling Technology, 80076S) | AKT (1:100; Cell Signaling Technology, 9272S) |
| V5–p-AKT(S473) | p-AKT(S473) (1:50, Cell Signaling Technology, 4051S) | V5-tag (1:100, Cell Signaling Technology, 13202S) |
| Dock7–Cdc42 | Dock7 (1:100; Santa Cruz Biotechnology, sc-398888) | Cdc42 (1:100, Cell Signaling Technology, 2466S) |
| V5–Cdc42 | V5-tag (1:100, Cell Signaling Technology, 80076S) | Cdc42 (1:100, Cell Signaling Technology, 2466S) |
